# *N*-glucosyltransferase GbNGT1 from *Ginkgo* complement auxin metabolic pathway

**DOI:** 10.1101/2020.08.30.249292

**Authors:** Qinggang Yin, Jing Zhang, Shuhui Wang, Jintang Cheng, Han Gao, Cong Guo, Lianbao Ma, Limin Sun, Shilin Chen, An Liu

## Abstract

As a group of the most important phytohormone, auxin homeostasis is regulated in a complex manner. Generally, auxin conjugations especially IAA glucosides are dominant on high auxin level conditions. Former terminal glucosylation researches mainly focus on *O*-position, while IAA-*N*-glucoside or IAA-Asp-*N*-glucoside has been neglected since their found in 2001. In our study, IAA-Asp-*N*-glucoside was firstly found specifically abundant (as high as 4.13 mg/g) in ginkgo seeds of 58 cultivars from Ginkgo Resource Nursery built in 1990. Furthermore, a novel *N*-glucosyltransferase GbNGT1, which could catalyze IAA-Asp and IAA to form their corresponding *N*-glucoside, was identified through differential transcriptome analysis and *in vitro* enzymatic test. The enzyme was demonstrated to possess specific catalyze capacity toward the *N*-position of IAA-amino acid or IAA among 52 substrates, and was typical of acid tolerance, metal ion independence and high temperature sensitivity. Docking and site-directed mutagenesis of this enzyme confirmed that E15G mutant could almost abolish enzyme catalytic activity towards IAA-Asp and IAA *in vitro* and *in vivo*. The IAA modification of GbNGT1 and GbGH3.5 was verified by transient expression assay in *Nicotiana benthamiana*. In conclusion, our results complement the terminal metabolic pathway of auxin, and the specific catalytic function of GbNGT1 towards IAA-amino acid provide a new way to biosynthesis indole-amide compounds.

**Highlight:** The N-glucosylation of IAA or IAA-amino acids in auxin metabolism had been neglected over decades, our work for GbNGT1 redeems the missing chain of auxin metabolic pathway.

## Introduction

The plant hormone auxin (indole-3-acetic acid, IAA) was discovered about 70 years ago. Our understanding of IAA signaling pathway has thrived during the last few decades, in contrast, key enzymes in its metabolic pathway were missing and the regulation of auxin metabolism was poorly understood (Ljung, 2013). In plant, IAA level could be attenuated by conjugation (mainly to amino acids and sugars) (Staswick *et al*., 2005); IAA conjugates are regarded as either reversible or irreversible storage compounds, although their functions and their regulation genes during plant growth and development is still under investigating (Korasick *et al*., 2013).

Amide-linked IAA conjugates (IAA-AA) constitute approximately 90% of the IAA pool in *Arabidopsis thaliana* (Tam *et al*., 2000). Many enzymes involved in IAA conjugation and IAA conjugate hydrolysis have been identified, such as auxin-inducible GRETCHEN HAGEN3 (GH3) family of amido synthases and different amido hydrolases (Staswick *et al*., 2005). Since glycosylation can alter many characteristics of aglycones in respect to their bioactivity, solubility, as well as their cellular localization, the procedure is considered as an important regulatory mechanism for cellular homeostasis and phytohormone activity (Ostrowski *et al*., 2014). Szerszen *et al*. (1994) firstly cloned an IAGlc synthase cDNA as coding an *O*-glycosyltransferase (OGT, IAGLU) from a maize library by antibodies (Szerszen *et al*., 1994). Subsequently, UGT84B1, UGT74E2, and UGT74D1 were identified sharing similar function with IAGlc synthase to form 1-*O*-IAA-glucoside in Arabidopsis (Jackson *et al*., 2001). Until Ljung *et al*. (2001) detected IAA-*N*-glucoside and IAA-Asp-*N*-glucoside in Scots pine (*Pinus sylvestris*), researchers started to notice the new metabolic branch for IAA (Ljung *et al*., 2001). However, no *N*-glucosyltransferase (NGT) gene was found up to now, even though another two new IAGlc synthases, OsIAGT1 and OsIAGLU were recently reported to produce IAA-*O*-glucoside in rice (*Oryza sativa*) (Liu *et al*., 2019; Yu *et al*., 2019).

Compared with *O*-glucosyltransferase (OGTs) reported in plant, NGTs were rarely identified, especially for small molecules and metabolites (Guo *et al*., 2015). IAA-*N*-glucoside was thought to be an irreversible form of IAA, for it’s harder to be hydrated by hydrolases than the IAA-*O*-glucoside (Casanova-Sáez *et al*., 2019). Known NGTs not only always possess substrate diversity, but also function as OGT, *S*-glucosyltransferase (SGT) or *C*-glucosyltransferase (CGT). AtUGT72B1 (*A. thaliana*), a NGT, was demonstrated to glucosylate toward the OH of 2, 4, 5-trichlorophenol, NH2 of 2, 3-dichloraaniline and SH of 4-chlorothiophenol (Brazier-Hicks *et al*., 2007). MiCGT (*Mangifera indica*) exhibited a robust capability of stereospecific *C*-glycosylation for 35 structural diverse drugs, such as scaffolds and simple phenols, using UDP-glucose as sugar donor. In the meanwhile, it could also form *O*- and *N*-glycosides (Chen *et al*., 2015). In medicinal plant, TcCGT1 (*Trollius chinensis*) was confirmed to catalyze *C*-, *O*-, *N*-, and *S*-glycosylation reactions(He *et al*., 2019). The substrates specificity of glycosyltransfeases mainly depend on their structures, rather than their preliminary sequences (Ostrowski *et al*., 2014). Combined with crystal structure of PtUGT1 (*Polygonum tinctorium*) which could glucosylate the OH of indoxyl (Hsu *et al*., 2018), the available structures of the above GTs could be used to explore the structural mechanism of NGTs for IAA.

Herein, we firstly found IAA-Asp-*N*-glucoside abundantly accumulated in ginkgo seeds, which was identified to inhibit cough (Liu *et al*., 2018). The content of IAA-Asp-*N*-glucoside in 58 ginkgo cultivars from China and Japan (two cultivars were introduced from Japan in 1990s) were among 1.02-4.13 mg/g D. W., significantly higher than that in rice seeds (∼ 0.03 *µ*g/g D. W.) (Kai *et al*., 2007). We found a unique GbNGT1 could catalyze *N*-position and format IAA-Asp-*N*-glucoside and IAA-*N*-glucoside through screening candidate GbUGTs from differential transcriptomes of ginkgo seeds and leaves. Using UDP-glucose as sugar donor, GbNGT1 specifically glucosylated the *N-*position of IAA-AAs or IAA among 52 substrates in enzymatic experiment. Docking analysis and site-directed mutagenesis experiments demonstrated that the E15 residue played critical role in *N*-glucosylating activity *in vitro and in vivo*. The *in vivo* function of GbNGT1 was confirmed by transient expression assay in *Nicotiana benthamiana*. These results not only improved the metabolic pathway of IAA, but also provided new tools for its protein engineering and biosynthetic research.

## Materials and methods

### Materials and chemicals

Ginkgo leaves, seed coats and seeds at different developing stages from June 15th to September 15th were collected in Beijing botanical garden; it is about 70 years old. Mature seeds of 58 cultivars were collected on October in Pizhou Resource Nursery. These samples were immediately frozen in liquid nitrogen, and stored at −80°C for further use. The substrates tested in the study were purchased from Xili Limited Co. (Shanghai, China) and Indofine (Hillsborough, NJ, USA). UDP-glucose, UDP-galactose and UDP-glucuronic acid were purchased from Sigma-Aldrich (Oakville, CA, USA). UDP-rhamnose was enzymatic synthesized using methods mentioned in Rautengarten et al. (2014) (Rautengarten *et al*., 2014). All chemicals used in this study were analytical or HPLC grade.

### Chemical synthesis of Substrates

Sixteen substrates were synthesized using chemical methods, including N-indole 3-acetyl-L-aspartic acid dimethyl ester (IAA-Asp (OMe)-OMe), N-indole 3-acetyl-L-aspartic acid (IAA-Asp), N-5-methylindole 3-acetyl-L-aspartic acid (IAA-Me-IAA-Asp), N-5-bromoindole 3-acetyl-L-aspartic acid (5-Br-IAA-Asp), N-indole 3-acetyl-L-glutamic acid dimethyl ester (IAA-Glu (OMe)-OMe), N-indole 3-acetyl-L-glutamic acid (IAA-Glu), N-5-methylindole 3-acetyl-L-glutamic acid (5-Me-IAA-Glu), N-5-bromoindole 3-acetyl-L-glutamic acid (5-Br-IAA-Glu), N-indole 3-acetyl-glycine (IAA-Gly), N-5-methylindole 3-acetyl-glycine (5-Me-IAA-Gly), N-5-bromoindole 3-acetyl-glycine (5-Br-IAA-Gly), N-indole 3-acetyl-leucine (IAA-Leu), N-5-methylindole 3-acetyl-leucine (5-Br-IAA-Leu). The synthesis steps and compound structures were confirmed by NMR as (Appendix S).

### Analysis of IAA-AA-*N*-glucosides metabolites by HPLC

50 mg dry weight sample was extracted with 2.5 ml methanol (25%) in an ultrasonic bath at 25°C for 30 min. Then the supernatant was filtered through a membrane (pore diameter is 0.22 *µ*m) after centrifugating (12, 000 rpm) for 10 min under 4°C. Finally, a 10 *µ*l aliquot was injected for subsequent analysis. The HPLC was used to determine the components based on the followed chromatographic separation terms with 280 nm or 220 nm detecting wavelength.

Chromatographic separation was achieved on a Venusil innoval C18 (250 mm × 4.6 mm, 5 *μ*m), with column temperature maintaining at 30 °C, auto-sampler temperature setting at 4 °C, using 0.1% formic acid in water and acetonitrile as solvent A and B. The injection volume per sample was 10 *μ*L and the flow rate was1.0 mL/min. Elution conditions were as follows: 0-30 min, B from 5% to 100%; 30-35 min, B from 100% to 100%.

### RNA-seq, candidate genes’ sequence analysis and gene clone

In order to examine the expression patterns of GbUGT and GbGH3s genes associated with IAA-*N*-glucoside and IAA-AA-glycosides biosynthetic pathway, RNAs from leaves and seeds of different developing stages were sequenced by Illumina HiSeq2000 platform. RNA extraction, sequencing and reads filtering were followed as previously described in Yin *et al*.,(2020) (Yin *et al*., 2020). Finally, thirteen GbUGT genes and eleven GbGH3s were selected according to the published genome and transcriptome of *G. biloba*. Multiple sequence alignment of deduced amino acid sequences was carried out by DNAMAN; predicted amino acid sequences of UGTs and GH3s were aligned using Clustal X2. Then, mixed cDNAs from leaves and seeds of *G. biloba* were used for gene amplification. For GbUGTs, the PCR products were purified and digested using the corresponding restriction enzymes, and then ligated to a pMAL-c2x vector (New England BioLabs, Ipswich, MA, USA) digested with the same restriction enzymes for expression of recombinant proteins in *Escherichia coli*. For *GbGH3s* and *AtGH3.6* genes, gateway system was used to construct pK7WG2D vector for verified the IAA modification in *planta*.

### Enzyme assay and products identification

Purification of recombinant UGT proteins in *E. coli* and enzymatic activity tests were done with minor modifications as previously described (Yin *et al*., 2017). In enzymatic test or kinetic analysis of the recombinant GbUGT proteins, purified enzymes (1-2 *µ*g) were incubated in reaction mixtures comprising 10 mM DTT, 50 mM Tris-HCl (pH 7.0), and 2 mM UDP-glucose, UDP-glucuronic acid, UDP-galactose, or UDP-rhamnose crude (20*µ*l enzymatic crude solution containing UDP-rhamnose), in a final volume of 50 *µ*l. The concentration of the tested acceptor substrates ranged from 100 to 2000 *µ*M. Reactions was stopped by adding methanol after 30 min’s incubation at 37°C. Samples were centrifuged at 14, 000 rpm for 10 min under 4°C, and further analyzed by HPLC using above mentioned procedure. The kinetic parameters *K*_m_ and *k*_cat_ were calculated by the Hyper 32 program (http://hyper32.software.informer.com/).

Enzymatic reaction solution was filtered through 0.22 *µ*m membranes, 10 *µ*L aliquot was used to analyze new products by HPLC, and then 1*µ*L aliquot was injected in UPLC-MS/MS to identify the products. UPLC-QTOF-MS/MS detection used 6540 Agilent 1260 photodiode array. Electrospray ionization (ESI) was applied in positive (PI) mode for MS and MS/MS to collect fragments information of the molecular weighs. The positive mode parameters were optimized as follows: HV voltage, 3.5 kV; capillary, 0.095 *μ*A; nozzle voltage, 1500 V; gas flow, 8 L min^-1^; gas temp, 320°C; nebulizer, 35 psi; sheath gas temp, 350°C; Sheath gas flow, 12 L min^-1^; and scan range at m/z 50-1250 units. Collision energy of 15V was used during MS/MS analysis.

### Homology modeling and docking statistic

Homology models of GbNGT1 were built, using the 3D structure of UGT72B1 (PDB No. 2VCH) and PtUGT1 (5NLM) as template, through SWISS-MODEL server at http://swissmodel.expasy.org. UDP-Glucose and IAA-Asp were respectively docking with the model structure of GbNGT1 using igemdock 2.1 program. The model with UDP-Glc and IAA-Asp was visualized by Pymol molecular graphics system at http://www.pymol.org.

### Functional Characterization of GbNGT1 in *N. benthamiana*

*GbNGT1*, mutant *E15G, AtGH3.6* (F24B18.13, At5g54510) and *GbGH3s* sequence were subcloned into the binary vector pK7WG2D by Gateway LR protocol. The *A. tumefaciens* GV3101 clone which contained *GbNGT1, GbGH3s, AtGH3.6* or *E15G* gene was incubated in 50 mL LB culture (containing 50 mg L^-1^ spectinomycin, 50 mg L^-1^ rifampicin) for overnight growth (200 rpm, 28°C). The culture was centrifuged for 10 min under 3000*×*g, and *A. tumefaciens* precipitation was resuspended and washed with infiltration buffer (10 mM MES, 10 mM MgCl_2_, and 100 *μ*M acetosyringone). The bacterial solution was adjusted to a final OD_600 nm_=0.5. The transiently transformation assay was conducted on 4-week *N. benthamiana* plants growing under 16/8 h light/dark rhythms thought leaf infiltration. LB 985 NightShade (Berthold Techonologies) and OLYMPUS IX73 were used to determine whether *GbNGT1, E15G, AtGH3.6* or *GFP* were transient expressed in tobacco leaves (Fig S2). After 24 h, the substrates IAA (4 mM) or IAA-Asp (4 mM) were infiltrated to the leaves of transformed above genes *N. benthamiana* respectively. Subsequently, the infected leaves were harvested, weighted (during 100-200 mg), grounded and then extracted with 500 *μ*L methanol solution (V_methanol_: V_water_=7:3). After sonicated for 30 minutes, the samples were centrifugated at 12, 000 rpm and filtered through membrane (pore diameter is 0.22 *µ*m). Finally, 10 *µ*l aliquot was applied for HPLC quantified analysis, and 1 *µ*l samples were used to UPLC-QTOF qualified analysis.

### Statistical analysis

Statistical analyses were performed by Excel (Microsoft Office, Microsoft). P-values were calculated using an unpaired, two-legged Student’s *t*test (**p < 0.01; *p < 0.05; ns, not significant). Data represent means ± standard deviation (n ≥ 3).

### Accession numbers

GenBank accession number MN908522 represents GbNGT1 or UGT717A21; while, MN908517 for GbGH3.5. The details of others candidate genes were listed in table S3.

## Results

### IAA-AA-*N*-glucosides concentrated in *G. biloba* seeds

IAA-AA-*N*-glucosides were confirmed to be pharmacological activity compounds in cough treatment in ginkgo seeds (Liu et al. 2018). NMR determined these compounds including IAA-Asp-*N*-glucoside and IAA-Glu-*N*-glucoside, which commonly distributed in mature seeds of 58 ginkgo cultivars or strains from Pizhou Resource Nursery built in 1990 (Supporting Information Figure S1&Table S1). The minimum content of IAA-Asp-*N*-glucoside is 1.02 mg/g D.W. in “Dalongyan, Anhui Quanjiao”, while the maximum content is up to 4.13 mg/g D.W. in “No. 18 Xincun”. The content of IAA-Glu-*N*-glucoside is less than that of IAA-Asp-*N*-glucoside, ranging from 0.24 to 1.23 mg/g D.W. in “No. 1-6, Hubei Anlu” and “No. 2, Guizhou Zhengan” respectively. It is worth mentioning that two cultivars from Japan also accumulated high content of IAA-AA-*N*-glucosides; the content of IAA-Asp-*N*-glucoside and IAA-Glu-*N*-glucoside in “Hisatoshi” or Jiushou” is 2.51 and 0.63 mg/g D. W., while in “Teng Kuo” or Tengjiulang” is 1.88 and 0.55 mg/g D.W, respectively.

Concurrently, we analyzed IAA-AA-*N*-glucosides content in different tissues of various developmental stages from June 15th to September 15th, including leaves, seed coats and seeds (Figure 1a). IAA-Asp-*N*-glucoside content in seeds (ranging from 1.5-2.0 mg/g D.W.) was at least 10-fold higher than that in seed coats (ranging from 0.07-0.20 mg/g D.W.) and leaves (ranging from 0.05-0.13 mg/g D.W.); whereas, IAA-Glu-*N*-glucoside content span in different tissues is smaller, in seeds that was 0.60-0.80 mg/g D.W., in seed coats was 0.04-0.06 mg/g D.W., and in leaves was 0.21-0.30 mg/g D.W. The IAA-Asp-*N*-glucoside content in seeds and leaves reached the peak in July, while in seed coats it gradually decreased from June to September. IAA-Glu-*N*-glucoside content of seeds, seed coats and leaves reached the maximum at July, June and August, respectively.

**Figure 1.**
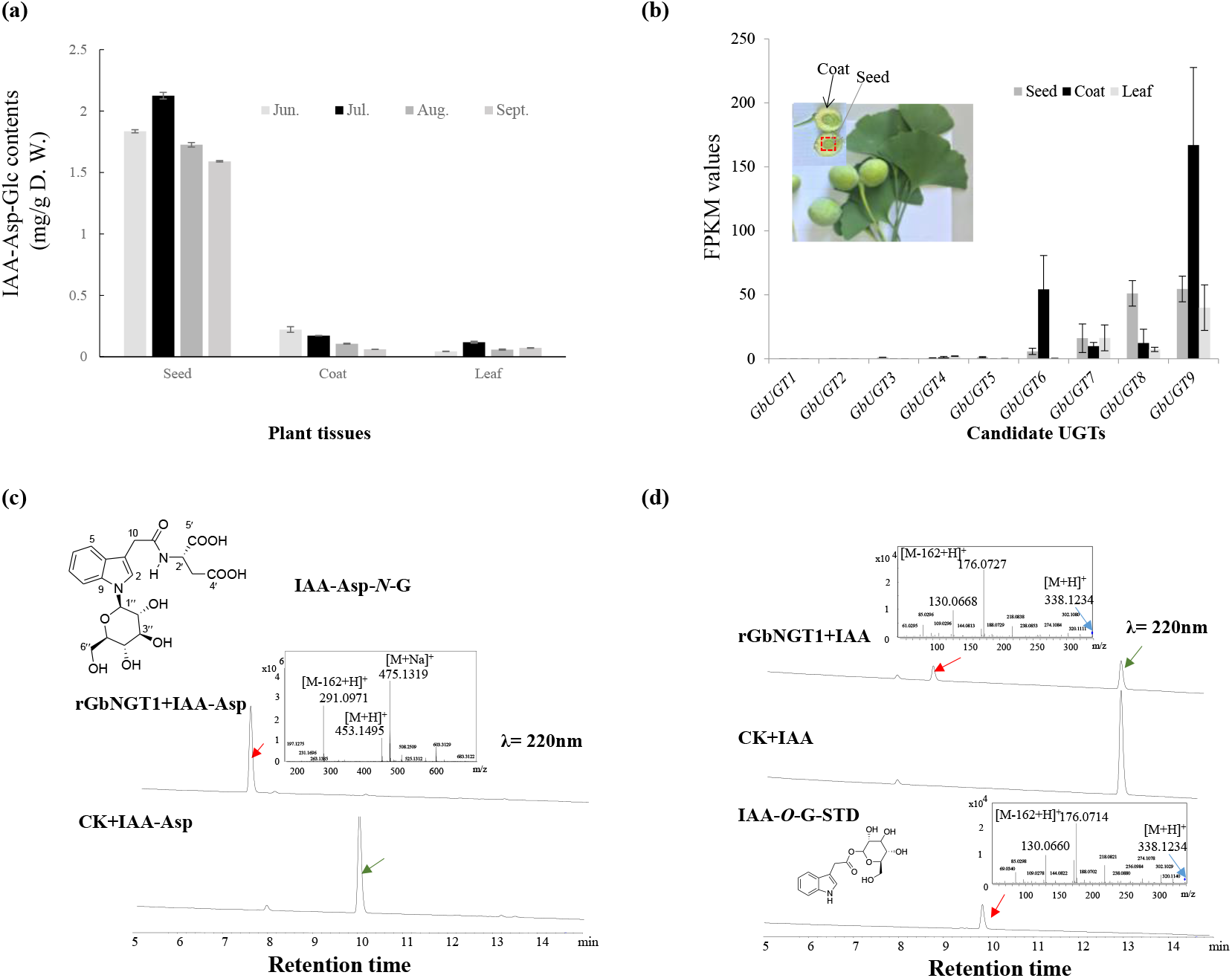
GbNGT1 catalyzed the formation of IAA-*N*-glucoside and IAA-Asp-*N*-glucoside. *(a)* The IAA-Asp-*N*-glucoside content in different tissues at the development stages. *(b)* The transcript levels of nine cloned GbUGTs in samples collected in June 15. *(c)* The new IAA-Asp-glucoside occurred in enzymatic products by HPLC and MS. *(d)* The new IAA-*N*-glucoside occurred in enzymatic products by HPLC and MS.

To be amazed, the presumed precursors IAA-AA or IAA-*N*-glucoside of IAA-AA-*N*-glucoside were extremely trace existed in all tested tissues; these compounds could only be qualitatively detected by UPLC-Q-TOF. The significant difference in contents of IAA-AA-*N*-glucosides and their precursors indicated that there must be some UGTs existed in ginkgo seeds which efficiently catalyzed the *N*-glucosylation of IAA-AA.

### Identification of ginkgo *N*-glucosyltransferase towards IAA and IAA-AA

Using Arabidopsis UGTs as queries, GbUGTs were screened from the available genome of *G. biloba* (Guan *et al*., 2016); concomitantly, final GbUGTs source was formed by UGTs containing the conserve domain PSPG (Plant Secondary Product Glycosyltransferase) boxes of plant UGTs. Combined with differential transcriptome analysis of *G. biloba*, 9 GbUGTs were cloned out of 13 candidates (Figure 1b, Supporting Information Table S2&3). In the prokaryotic expression experiment (Supporting Information Figure S2), GbUGT8 recombinant protein was identified to be able to catalyze IAA-Asp to produce a new product using UDP-glucose as sugar donor. Comparing to the substrate, the new product gained 162 molecular weight detected by Mass spectrometry and was further identified as IAA-Asp-*N*-glucoside by NMR analysis (Figure 1c & Supporting Information Table S4). However, GbUGT8 could not catalyze IAA-Asp to produce new products using UDP-galactose, UDP-glucuronic acid or UDP-rhamnose as sugar donor. Similar to Asm25 (*Actinosynnema pretiosum*), which catalyzes *in vitro* glycation of PNDs at the macrolactam amide nitrogen position using UDP-glucose as the sole sugar donor (Zhao *et al*., 2008), GbUGT8 could only use UDP-glucose as sugar donor to catalyze *N*-glucoslation of IAA-Asp. Then it was finally named as GbNGT1.

Meanwhile, it was found that the recombinant protein also could catalyze the *N*-glucosylation of IAA (Figure 1d). The mass spectrum showed that the new product was IAA glucoside, but not IAA-*O*-glucoside judging by retention time of the standard. In the meantime, the MS product fragments did not contain 218.1234 [M+H-120]^+^ and 248.1234 [M+H-90]^+^ from parent ion 338.1234 [M+H] ^+^of IAA glucoside, which were the characterize fragments of *C*-glucosides (Chen *et al*., 2015). Therefore, the product was predicted to be IAA-*N*-glucoside. As to notify, this product was also commonly found in seeds of *G. biloba* (Supporting Information Figure S3). It indicated that IAA-*N*-glucoside could be produced through the *N*-glucosylation of IAA in plant.

### The specificity of GbNGT1

The enzymatic ability of GbNGT1 under pH 5.0 (138.55 nkat/mg protein), 6.0 (133.56 nkat/mg protein) and 7.0 (129.83 nkat/mg protein) for IAA-Asp were obvious higher than that under pH 8.0 (71.92 nkat/mg protein), indicated that it could tolerant acid (Supporting Information Figure S4). Significant differences of enzymatic ability were not found between at 25°C (81.37 nkat/mg protein) and 35°C (80.06 nkat/mg protein), whereas the catalyzed activities at 45°C (0.92 nkat/mg protein) and 55°C (0.37 nkat/mg protein) were severely weakened (Supporting Information Figure S); it indicated that GbNGT1 was sensitive to high temperature. In buffers with different metal ions, the enzymatic activity did not change significantly; the result showed the vitality of GbNGT1 was independent to metal ions (Supporting Information Figure S4).

Fifty-two substrates were tested for exploring whether GbNGT1 possessed substrate diversity the same as UGT72B1, MiCGT and TcCGT1; only three IAA-AAs could be glucosylated to form their corresponding IAA-AA-*N*-glucosides by MS and NMR (Supporting Information Figure S5 and Table S5&6), the converse rates were 92.3%, 23.7% and 80.5% towards IAA-Glu, IAA-Gly and IAA-Leu, respectively. In order to investigate the relationship between substrate electron density and enzymatic activity, we synthesized various kinds of IAA and IAA-AA derivatives as substrates, which incorporated strong electron donor moiety or electron absorption capacity (Supporting Information Appendix). Towards IAA-AAs adding electron donor or absorption moieties to the benzene ring, GbNGT1 directly lost its catalytic activity (Figure 2a). Similarly, the enzyme could not catalyze modified IAA when the electron donor or electron acceptor group was added. These results demonstrated that the activity of GbNGT1 was sensitive to electron and very likely strictly depend on steric stabilization.

**Figure 2.**
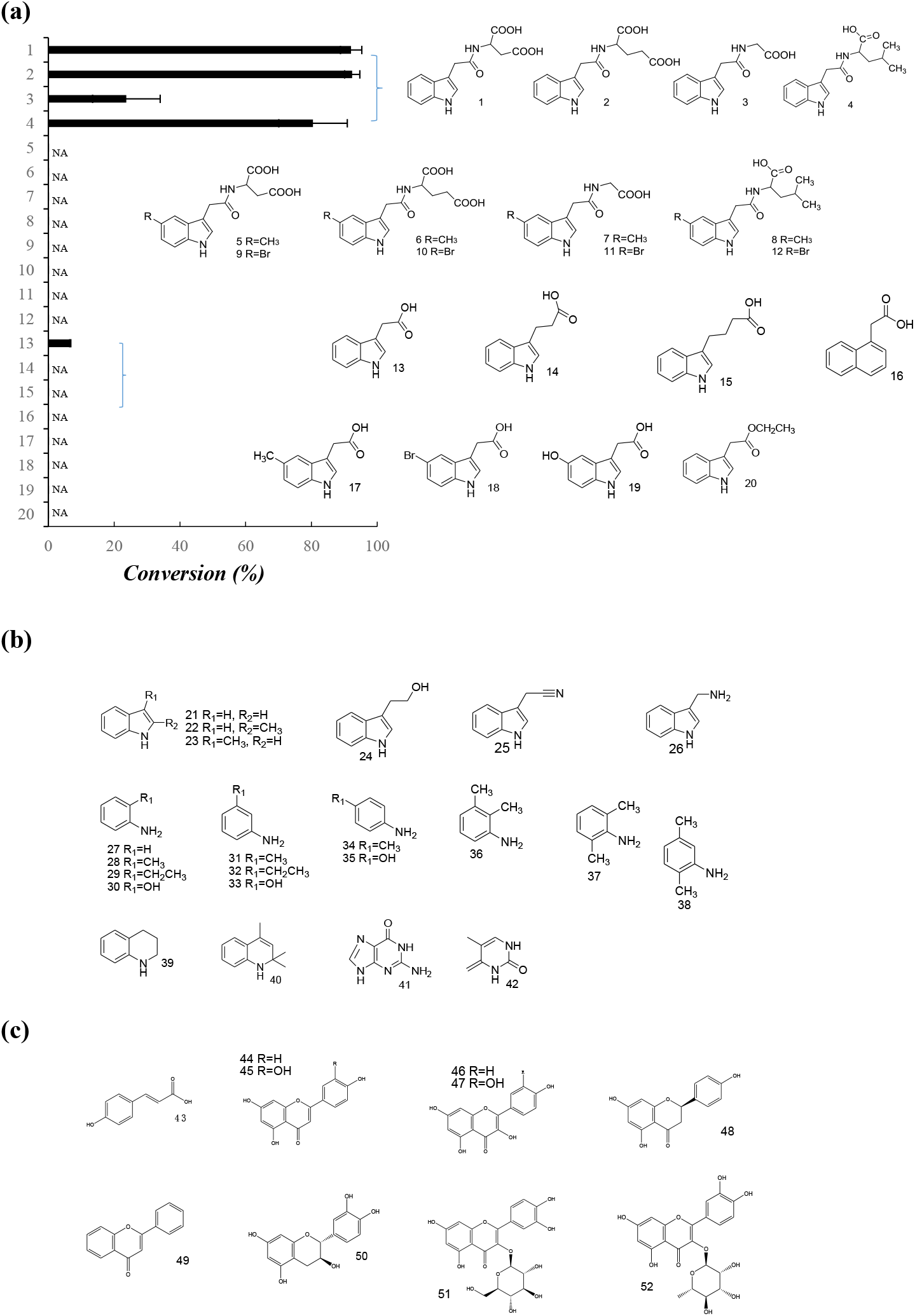
The enzymatic specificity of GbNGT1 as *N*-glucosyltranferase. *(a)* The conversion rates of GbNGT1 toward IAA-Asp, IAA-Asp derivatives, IAA, IAA analogues, and IAA derivatives. NA means no product detected. *(b)* The list of indole amide or anilines which could not be glycosylated by GbNGT1. *(c)* The flavonoids list which could not be glycosylated by GbNGT1.

Except for IAA and IAA-AA derivatives, 25 indole or aniline derivatives were used to test the universal activity of GbNGT1. It was found that GbNGT1 could not glucosylate IAA analogues, such as IBA, IPA and NAA. Furthermore, GbNGT1 could not catalyze aniline or indole derivatives to form new glycosylation products (Figure 2b), whatever the substitutes (-OH, methyl or ethyl) were nearby NH2 or N. These results suggested that GbNGT1 was a specificity *N*-glucosyltransfease, which strictly defined the substrate structure, tiny modification or changes of substrate could cause steric hindrance thereby directly affected the affinity between enzyme and substrate.

Ginkgo contains various kinds and a great number of flavonoid glucosides, thus it is reasonable to test whether GbNGT1 could catalyze flavonoids. 10 flavonoids including flavone, flavonol and proanthocyanidin monomers were used as substrates (Figure 2c); however, GbNGT1 could not catalyze any of the tested flavonoids. These results also indicated that GbNGT1 could not glucosylate the common *C*-position or *O*-position of substrates like other NGTs.

### The 15th residue Glu (E) determined the *N*-glucosylation function of GbNGT1

Based on crystal structures of UGT72B1 (PDB No., 2VCH) glycosylated toward *N*-, *S*- and *O*-position and PtUGT1 (5NLM) glucosylated at the OH of indoxyl (Brazier-Hicks *et al*., 2007; Hsu *et al*., 2018), we simulated the protein models of GbNGT1; obtained two model structures similarity between UGT72B1 and PtUGT1 were 31% and 30%, correspondingly. The former was selected to imitate molecular docking with UDP-glucose and IAA or IAA-AAs (Supporting Information Table S7). Among the 5 small molecules, the totally needed energy for IAA was the highest; it suggested that the enzymatic activity toward IAA would be lower comparing to IAA-AAs, which was coincided with our above experimental results (Figure 2a).

Docking results of IAA-Asp, UDP-glucose and protein model showed that there were eight key residues for binding substrates: His (H381), Trp (W384), Asn (N385) and Gln (Q386) were related to UDP-glucose by Hydrogen bond, while Glu (E15), Gln (Q16), Gly (G17) and Gly (G383) bonded to IAA-Asp by van der Waals interactions (Figure 3a & Supporting Information Fig S6). Alignments among GbNGT1, UGT72B1 and PtUGT1 showed the above mentioned residues were identical in position except E15 and G16 (Supporting Information Figure S6).

**Figure 3.**
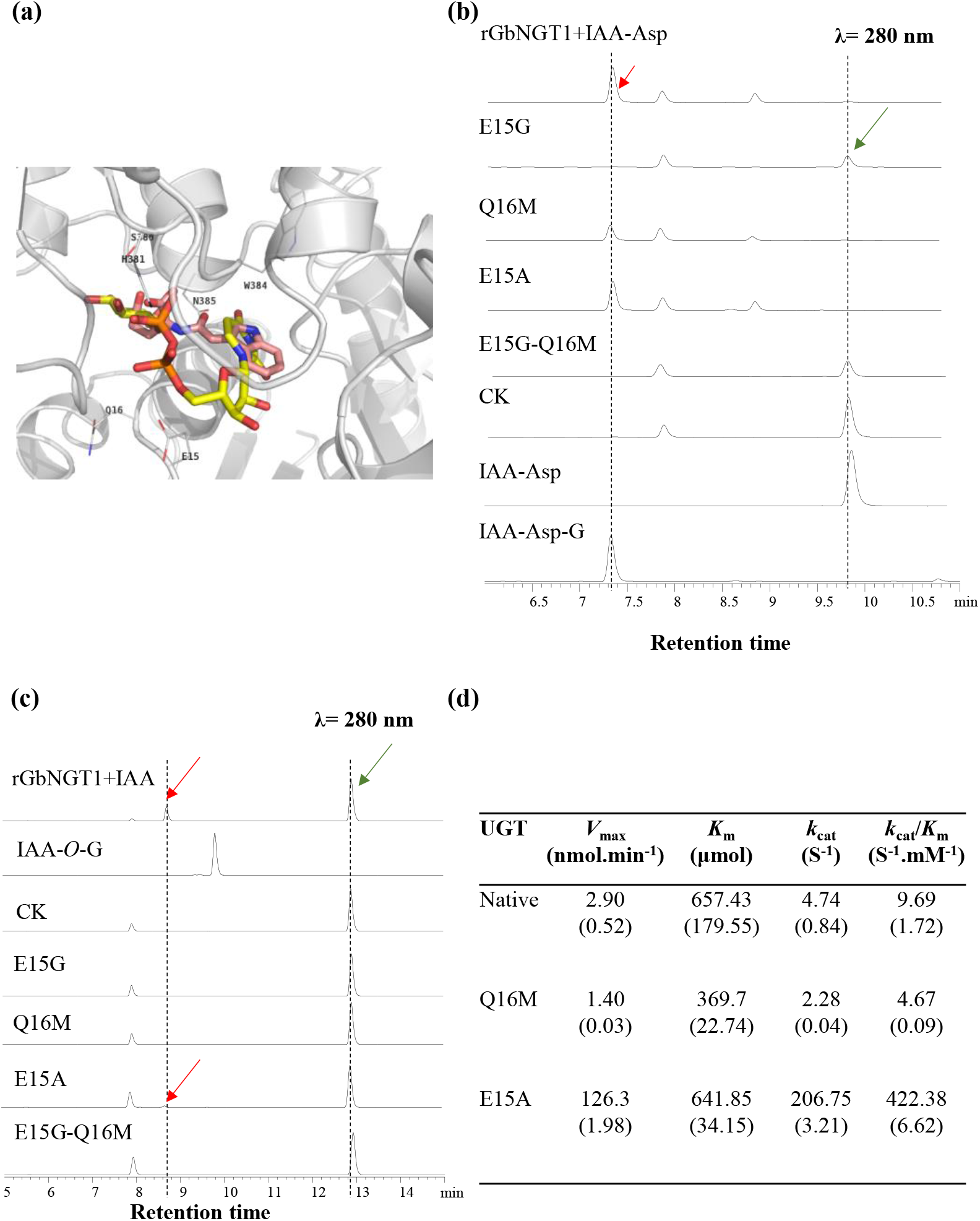
E15 determined the catalyzed activity of GbNGT1. *(a)* the binding domain of GbNGT1 docking with UDPG and IAA-Asp, the molecule marked yellow and orange is UDPG, that marked pink is IAA-Asp. *(b)* HPLC chromatograms of enzymatic products, which including native or mutants, UDPG and IAA-Asp. *(c)* HPLC chromatograms of enzymatic products, which including native or mutants, UDPG and IAA. *(d)* The kinetic data of native and mutants towards IAA-Asp, Averages (SD), n=3. Red arrow, product; green arrow, substrate.

Four GbNGT1 mutants were obtained by site-directed mutagenesis (Supporting Information Figure S7), including E15G, Q16M, E15A and double mutant of E15 and Q16, E15G-Q16M. Enzymatic tests showed that the activity of E15G and E15G-Q16M were abolished toward IAA-Asp; while E15A and Q16M still possess different catalyzed ability toward IAA-Asp (Figure 3b&d). E15A enzymatic efficiency towards IAA-Asp was dramatically increased comparing to native protein, with the *k*_cat_/*K*_m_ value significantly changing from 9.69 S^-1^ mM^-1^ to 422.38 S^-1^ mM^-1^. On the contrary, the *k*_cat_/*K*_m_ value of Q16M towards IAA-Asp was decreased by 4.67 S^-1^ mM^-1^. These results clarified that the combination of GbNGT1 and its substrates was sensitive to electron circumstance and steric distribution, which coincided with the conclusion of GbNGT1 enzymatic specificity.

Towards IAA, the enzymatic activities of mutants showed similar tendency with IAA-Asp; none glucosylation product was detected in the enzymatic experiments with E15G, Q16M or E15G-Q16M toward IAA. It is worth notifying that Ala replaced Glu in E15A could enhance the enzymatic efficiency towards IAA-Asp, but conversely reduced it towards IAA (Figure 3c). These results designated that E15 site of GbNGT1 determined its *N*-glucosylation towards IAA and IAA-Asp.

### The *in vivo* function of GbNGT1 in *N. benthamiana*

The *in vivo* function of GbNGT1 functionality was tested by *Agrobacterium tumefaciens*-mediated transient expression assay in *N. benthamiana* (Figure 4a, Supporting Information Figure S2). GbNGT1 coding sequences was cloned under the 35S promoter for the constitutive expression in *N. benthamiana*. Not surprisingly, IAA, IAA-Asp and IAA-Asp-*N*-glucoside were not detected in transgenic or wild type tobaccos by HPLC.

**Figure 4.**
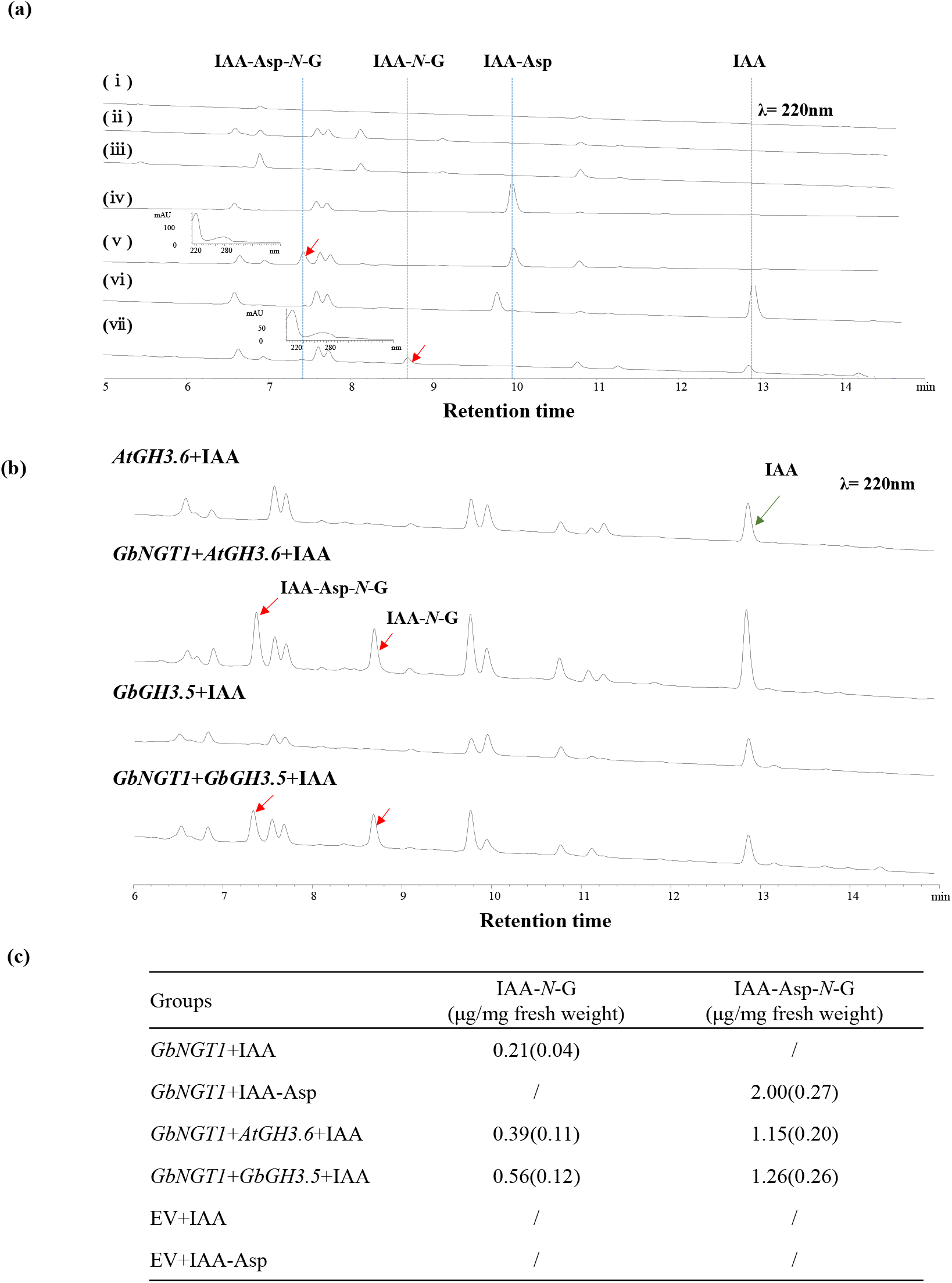
Functional Characterization of GbNGT1 in *N. benthamiana. (a)* The HPLC spectrums of *N. benthamiana* leaves with or without *GbNGT1*, wildtype and empty vector (EV): (i) wild type; (ii) empty vector-transformed leaves; (iii) *GbNGT1*-transformed leaves, (iv),empty vector -transformed leaves adding IAA-Asp, (v), *GbNGT1*-transformed leaves adding IAA-Asp, insert picture is the UV spectrum of product, (vi),empty vector-transformed leaves adding IAA; (vii) *GbNGT1*-transformed leaves adding IAA, insert picture is the UV spectrum of product. *(b)* GH3s and GBNGT1 reconstructed the IAA-Asp-N-Glucoside formation in tobacc. *(c)* the contents of IAA-*N*-glucosides in tobacco leaves transient expressed different genes combination, “/” means no product detected by HPLC, Averages (SD), n=3.

Therefore, IAA or IAA-Asp substrates were fed to the tobacco leaves. The corresponding products IAA-*N*-glucoside and IAA-Asp-*N*-glucoside could be found in *GbNGT1*-transformed tobacco leaves, but could not be detected in the control. The loss of function of mutant E15G was also verified in *N. benthamiana*, whatever towards IAA or IAA-Asp (Supporting Information Figure S7), accord with the enzymatic results.

For reconstructing the IAA modification in tobacco, *AtGH3.6* were introduced as it catalyzed the formation of IAA-Asp from IAA (Staswick *et al*., 2005). Expectedly, both IAA-*N*-glucoside and IAA-Asp-*N*-glucoside could be detected in *AtGH3.6*- and *GbNGT1*-transformed leaves adding IAA. Meanwhile, 11 candidates GbGH3s genes were transient expressed in tobacco with GbNGT1, only *GbGH3.5* instead of *AtGH3.6*, above two IAA *N*-glucosides were detected in *GbGH3.5*- and *GbNGT1*-transformed leaves (Figure 4b&c). Interestingly, IAA-*O*-glucoside was found in the control after adding IAA, while only IAA-*N*-glucoside was detected in *GbNGT1*-transformed leaves. In a word, the identification of GbNGT1 improves IAA metabolic pathway by directly filling the gap of the *N*-glucosylation of IAA or IAA-AA (Figure 5).

**Figure 5.**
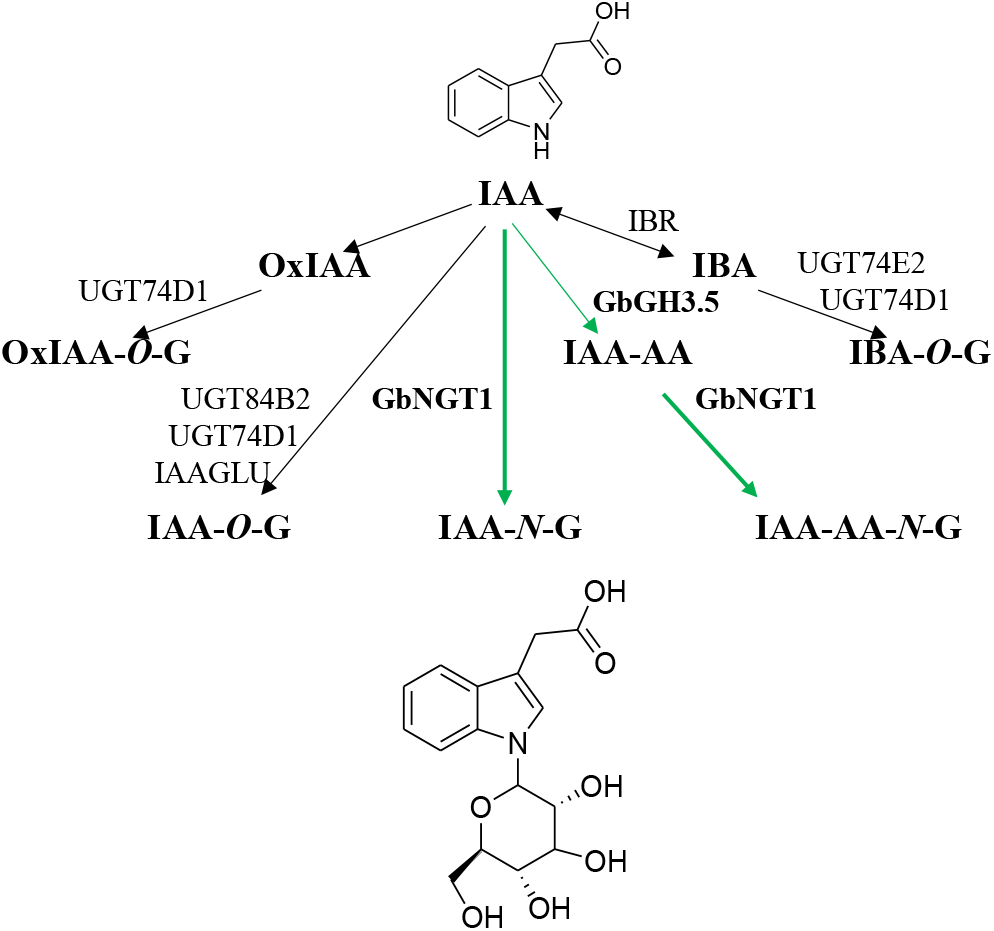
The renewed IAA metabolism pathway. The character G in IAA-*O*-G or IAA-Asp-*N*-G means glucoside.

## Discussion

### Discovery of IAA-Asp-*N*-glucoside enriching in ginkgo seeds promotes the supplement of IAA metabolic pathway

IAA is generally found trace amount in plants although in various styles, including IAA-esters (such as, IAA-*O*-glucoside) and IAA-AAs (Tam *et al*., 2000). The seeds of maize contained 79.5 *µ*g/g D.W. total IAA which was the highest one ever reported in literatures; while it was just 8 *µ*g/g D.W. in oat seeds, and less than 1 *µ*g/g D.W. in other tested plants (Bandurski *et al*., 1977). Herein, we found amazing amount of IAA-Asp-*N*-glucoside existed in ginkgo seeds. During 58 ginkgo cultivars, the IAA-Asp-*N*-glucoside content was up to 4 mg/g D.W., over thousand times comparing to that ∼0.15 *µ*g/g D.W. in rice seeds, the only plant which tested the content of this compound in published report (Kai *et al*., 2007). Meanwhile, slight IAA-*N*-glucoside was also detected by UPLC-QTOF in ginkgo seeds. The biosynthetic pathway of IAA-Asp-*N*-glucoside or IAA-*N*-glucoside, also called IAA *N*-glucosylation pathway, was totally unidentified due to the trace amount and few species that contain them, ever if these compounds were found in Scots pine 20 years ago (Ljung *et al*., 2001).

In this work, we found out IAA-Asp-*N*-glucoside commonly hyper-accumulating in ginkgo seeds through large scale resource screening. After differential transcriptome analysis, we firstly cloned and systematically identified specificity IAA *N*-glucosyltransferase (NGT) which completed the metabolic network of IAA. Based on the GbNGT1 enzyme activity toward IAA or IAA-Asp and the widely studied IAA-*O*-glucoside pathway, the formation of IAA *N*-glycoside may through two ways: (i) Amino acid is firstly added to IAA to form IAA-AA by GH3s; then it is glucosylated to IAA-Asp-*N*-glucoside by UGTs. Alternatively, (ii) IAA-*N*-glucoside is firstly produced by UGTs glucosylating IAA *N*-position; after that, GH3s catalyze it to format IAA-Asp-*N*-glucoside (Figure5). The result of transient expressed *GbNGT1* and *GbGH3.5* in tobacco have provide the possibility exited of first status, however, there may be differentiation or diversification of GH3s in ginkgo needing more GH3s function exploration.

The known metabolic pathway of IAA includes *O*-glucosyltransferases and amino acid conjugate synthetases. The OGTs and GH3s directly affect the existence form and dynamic equilibrium of IAA in plants; therefore regulate plant growth and development. The first reported IAA *O*-glucosyltransferase is IAGLU from maize identified in 1994 (Szerszen *et al*., 1994); subsequently, this kind of OGTs in Arabidopsis, duckweed, cauliflower, soybean, tomato, rice and tobacco were identified. They widely affected the structure, growth and development of plants, such as leaf angle and structure, dwarf, flower development via regulating the balance of IAA in plant (Ostrowski *et al*., 2014). Along with the identification of GH3 family which catalyzed the formation of IAA-amino acid in Arabidopsis, the GH3s in rice, moss, pea (*Pisum sativum*) and strawberry were reported to involve in anti-pathogen, shoot cell elongation, lateral root development and geotropism of root (Ding *et al*., 2008; Ostrowski *et al*., 2016; Tang *et al*., 2019). As a powerful enzyme, GbNGT1 not only determines the new IAA metabolic branch, but also plays an important role in downstream modification of the known IAA-AA branch. The identification of GbNGT1 largely complements the metabolic network of IAA; opens up new directions for homeostasis study of IAA in plant; and finally, must promotes the discovery of new IAA regulation models in medicinal plant.

As the main product of IAA amino acid conjugates, IAA-Asp was considered to be the irreversible form of IAA and no use in plant till its hydratases were found (Ljung K, 2013). In Chinese cabbage (*Brassica rapa*) the enzymatic activity of IAA–Asp hydrolysis increased in *Plasmodiophora brassicae*-infected root galls compared with control roots (Ludwig-Müller *et al*., 1996); in *M. truncatula*, MtIAR31, -32, -33, and -34 had hydrolytic activity against IAA-Asp and IBA-Ala (Campanella *et al*., 2008). At present, IAA-*N*-glucoside was also considered as a more stable pattern comparing to IAA-*O*-glucoside (Casanova-Sáez *et al*., 2019). However, such high content of IAA-Asp-*N*-glucosides concentrates in ginkgo seeds, what is the reason? And what kind of physiological function do they act? Are there any special hydratases which could cleave C-N bond to free IAA in ginkgo? Those questions should be paid more attention to in the future.

### Specificity of *N*-glucosylation

*N*-glycosylation modification in proteins or peptides is more common than in small molecules. Asparagine (Asn) was generally considered as the main amino acid site for *N*-glycosylation in peptides; this modification of Asn is characterized by covalent attachment of an oligosaccharide to its side chain amide (Chung *et al*., 2017). GbNGT1 could glucosylate small molecules including IAA-Asp, IAA-Glu, IAA-Leu, or IAA-Gly at the *N*-position of IAA residue, but not at the amide nitrogen of these amino acid residues’ side chain. Our results confirm the comment that the catalytic mode of *N*-glycosyltransferses towards small molecules and proteins are different (Naegeli *et al*., 2014).

IAA *N*-glucosides existed in monocotyledons (rice and maize) and dicotyledons (Arabidopsis, *Lotus japonicus*,) (Kai *et al*., 2007), gymnosperms (Scots pine and ginkgo) (Ljung *et al*., 2001) and even in microbes (*Cortinarius brunneu*) (Teichert *et al*., 2008); it means that the enzymes contributed to biosynthesis these compounds should be conserved. The homologous genes of OGTs for IAA or NGTs for other small molecules could be easily found in different species. It suggested that the UGT amino acid sequences are related to their function in plants. OsIAGLU showed 67% amino acid identity with ZmIAGLU, which possessed the same function to glucosylate the hydroxyl of IAA (Yu et al., 2019). Similarly, BnUGT1 and AtUGT72B1 shared 85% sequence identity and the same *O*-glycosylation function toward 3,4-dichlorophenol; differentially, only AtUGT72B1 could glucosylate the *N*-position of 3,4-dichloraaniline (Brazier-Hicks *et al*., 2007). Unusually, the highest similarity among GbNGT1 and other UGTs is just 44% from *Picea sitchensis* (GenBank: ABR17691.1) when blasted in NCBI database; moreover, the homologous genes of GbNGT1 could not be found by blasting the default value even in ginkgo genome. These results suggest that the catalytic capacity of *N*-glucosyltransferse toward IAA or IAA-AA may less dependent on amino acid sequences but mainly rely on protein structure.

The reported *N*-glycosyltransferases usually catalyze the different type glycosylation reactions at diversity sites. AtUGT72B1 could conduct the *O*-glycosylation of 2,4,5-trichlorophenol, *N*-glycosylation of 2,3-dichloraaniline and *S*-glycosylation of 4-chlorothiophenol (Brazier-Hicks *et al*., 2007). MiCGT could glycosylate the *C*-position of maclurin, *O*-position of phenol and *N*-position of 3,4-dichloraaniline (Chen *et al*., 2015). TcCGT1 was identified to catalyze four types of glycosylation, the *C*- or *O*-position of phenols and flavonoids, the *N*-position of 3,4-dichloraaniline, and *S*-position of 3,4-chlorothiophenol (He *et al*., 2019). In contrast, GbNGT1 specifically catalyzes the *N*-glucosylation toward IAA or IAA-AA.

Generally speaking, the relatively reserved PSPG motif located in C terminal of UGTs was bond to sugar donor, while the diversity residues close to N terminal were bond to sugar acceptor, which determined the specific selection of substrates (Ostrowski *et al*., 2014). So far, very few key residues in N terminal were identified even if some crystal structures of NGTs were revealed. H24 and E396 of TcCGT1 were demonstrated to play key role in the stabilization and location of small molecule substrates; in addition, the mutants I94E and G284K of TcCGT1 could convert enzymatic activity from *C*-to *O*-position (He *et al*., 2019). After converting all five amino acid residues within 314-320 of BnUGT1 to the corresponding key amino acids in UGT72B1, BnUGT1 gained the function of *N*-glycosylation activity toward 2,3-dichloraaniline. The *O*-glycosylation activity of UGT72B1 mutant H19Q was severely decreased toward 3,4-dichlorophenol, with *k*_cat_ value reducing to 1/300 of that in native protein. Simultaneously, the *N*-glycosylation capacity of H19Q toward 3,4-dichloraaniline was also dropped to 1/2 of that in native protein (Brazier-Hicks *et al*., 2007). In our study, amino acid alignments revealed H18 nearby N terminal of GbNGT1 was corresponding to the critical sites H19 in AtUGT72B1 and H24 in TcCGT1. Docking analysis identified another special residue E15 binding to IAA-Asp in GbNGT1. Mutating E15 (Glutamic acid) to E15G (Glycine) indeed drastically decreased or even abolished the catalytic activity of *N*-glycosylation toward IAA-Asp or IAA *in vitro* and *in vivo*; whereas, converting E15 to E15A (Alanine) oppositely enhanced enzymatic efficiency by forty-fold comparing to native protein toward IAA-Asp. E15 is another N terminal critical residue determining the specificity of substrates besides His in N-terminal of GbNGT1, which provides a new reference site for reconstruction of NGTs.

### The application prospect of GbNGT1

Ginkgo flavonoids and ginkgolides had been widely used as important drug and dietary supplements in the world (Su *et al*., 2017). IAA-Asp-*N*-glucoside was mentioned as the main pharmacological compound in ginkgo for inhibiting cough, antiasthmatic and eliminating phlegm (Liu *et al*., 2018);furthermore, there were considerable pharmacological studies on IAA derivates or indole alkaloids, for instance, IAA induced cell death in combination with UV-B irradiation by increasing apoptosis in PC-3 prostate cancer cells (Kim *et al*., 2010); indole-*N*-glucosides have the potential to serve as a novel SGLT2-selective inhibitors (Sodium-Glucose cotransporter) to cure type II diabetes (Nomura *et al*., 2013). It indeed suggests that IAA derivates own pharmaceutical developing prospect. GbNGT1 not only has high specific activity toward IAA-AA, but also possesses spatial specificity; furthermore, IAA-Asp-N-glucoside was produced from IAA in tobacco after transforming *GbGH3.5* and *GbNGT1*; it indicated that these proteins are potential engineering enzymes for the biosynthesis of IAA-AA-*N*-glucoside.

## Conclusions

The *N*-glucosylation of IAA or IAA-amino acids in auxin metabolism had been neglected over decades, our work for GbNGT1 redeems the missing chain of auxin metabolic pathway. The unique *N*-glucosylation function and high efficiency of GbNGT1 in IAA metabolic pathway distinguish the uncommon protein from other published *N*-glycosyltransferases with multiple functions; the function determined crucial residue (E15) nearby N terminal inspires new way exploring for glucosyltransferase modification. The surprising abundant accumulation of IAA metabolites in ginkgo seeds promotes the discovery of this neglected branch; it sets up a good example for enriching major metabolic studying in special medicinal plants.

## Supporting information

total supplemental information

## Supplementary data

Figures S1. The IAA-AA-N-glucosides content of different cultivars. (a) The IAA-Asp-N-glucoside content in seeds of 58 cultivars. (b) The IAA-Glu-N-glucoside content in seeds of 58 cultivars.

Figures S2. IAA-N-glucoside existed in ginkgo seeds. (a) MS spectrums extracted 338.1234 from samples of enzymatic product, ginkgo seeds and IAA-O-glucoside. (b) The MS spectrum related to chart A.

Figures S3. GbNGT1 expressed in E.coli and N. benthamiana. (a) SDS-PAGE gel of recombinant GbNGT1 protein; (b) and (c) transient expressed GFP or GbNGT1-GFP in tobacco was confirmed by LB 985 NightShade (Berthold Techonologies) and OLYMPUS IX73, respectively, WT, wildtype, EV, empty vector.

Figures S4. The enzymatic activity of GbNGT1 toward IAA-Asp substrate in solution at different pH value (a), temperature (b) and metal ion (c).

Figures S5. HPLC and MS spectrums of GbNGT1 with IAA-Glu (a, S2), IAA-Gly (b, S3), and IAA-Leu (c, S4). Negative ion PI model was used to detect the three substrates, S2a, S3a and S4a are the new products in enzymatic reactions.

Figures S6. The key residues predicted by docking. (a) Amino acid alignment of GbNGT1, AtUGT72B1 and PtUGT1, asterisks for binging amino acids. (b) The overall chart of GbNGT1 docking with UDPG and IAA-Asp, the molecule marked yellow and orange is UDPG that marked pink is IAA-Asp.

Figures S7. The functions of GbNGT1 mutants in *E. coli* and *N. benthamiana*. (a) The SDS-PAGE gel of native and mutant recombinant proteins of GbNGT1. M, maker; CK, empty vector. (b) The HPLC spectrums of N. benthamiana leaves transformed mutant E15G or GbNGT1: (i) mutant E15G-transformed leaves adding IAA-Asp; (ii) GbNGT1-transformed leaves adding IAA-Asp; (iii) mutant E15G-transformed leaves adding IAA; (iv), GbNGT1-transformed leaves adding IAA. Red arrow, new product; green arrow, substrate.

Tables S1. The 58 ginkgo cultivars used in our study.

Tables S2. The transcripts of 13 UGTs and 11 GbGH3s from the public data. I-fruit, immature fruit, R-fruit, ripe fruit.

Tables S3.The transcripts of cloned UGTs and GH3s according to transcriptome data collected on June 15th, 2018

Tables S4. 1H-NMR and 13C-NMR spectrum data of IAA-Asp-N-Glc compounds

Tables S5. 1H-NMR and 13C-NMR spectrum data of IAA-Gly-N-Glc compounds

Tables S6. 1H-NMR and 13C-NMR spectrum data of IAA-Leu-N-Glc compounds

Tables S7. The predicted docking energy of GbNGT1 with IAA and IAA-AAs. The output data included total energy (Kcal/mol), van der Waals interactions (VDW, Kcal/mol), Hydrogen bonding (HBond, Kcal/mol), electrostatic interactions (Elec Kcal/mol), and average conpair (AverConPair).

Appendix S. Synthesis of Substrates and NMR information for used compounds.

## Acknowledgments

We thank Dr. Xiaoyan Han (Institute of Botany, Chinese Academy of Sciences) for key suggestions and subtle corrections about the paper. This research was supported by Beijing Natural Science Foundation of China (7192138), the National Natural Science Foundation of China (81703647), the Fundamental Research Funds for the Central public welfare research institutes of China (ZZ13-YQ-097), and National Key R&D Program of China (2019YFC1711100).

## Author Contributions

QY and AL designed research; QY, JZ, SW, JC, HG, and CG performed research; QY, AL and LM collected samples; QY, JZ, and HG analyzed data; and QY wrote the paper; SC, and AL revised the paper

