## Supplementary material for "*N*-glucosyltransferase GbNGT1 from *Ginkgo* complement auxin metabolic pathway": total supplemental information

**This PDF file includes:**

Figures S1 to S7

Tables S1 to S7

Appendix S

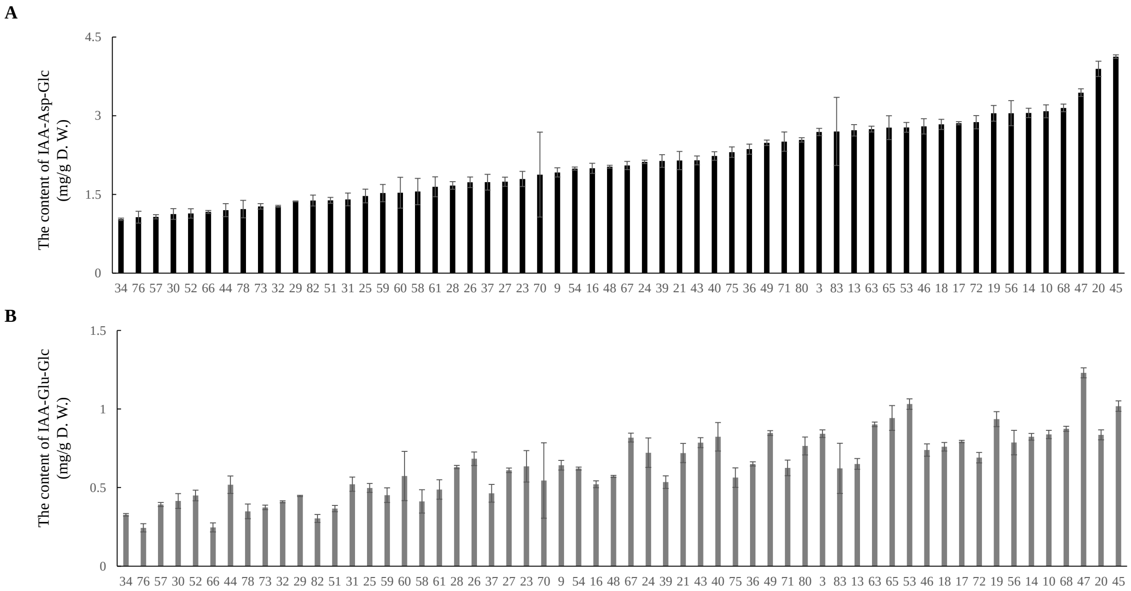


Fig. S1. The IAA-AA-*N*-glucosides content of different cultivars. *(A)* The IAA-Asp-*N*-glucoside content in seeds of 58 cultivars. *(B)* The IAA-Glu-*N*-glucoside content in seeds of 58 cultivars.

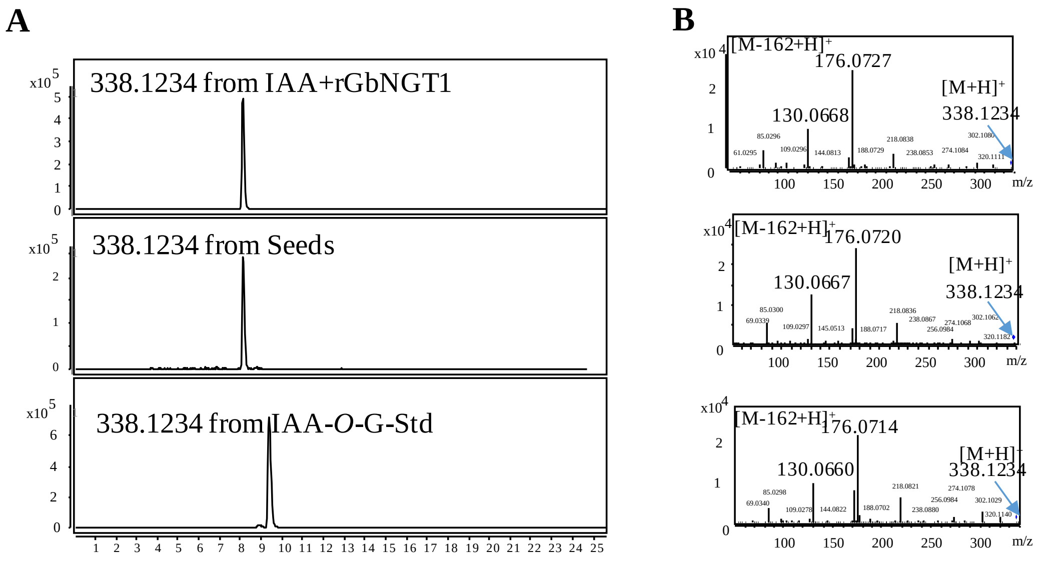


Fig. S2. IAA-*N*-glucoside existed in ginkgo seeds. *(A)* MS spectrums extracted 338.1234 from samples of enzymatic product, ginkgo seeds and IAA-*O*-glucoside. *(B)* The MS spectrum related to chart A.

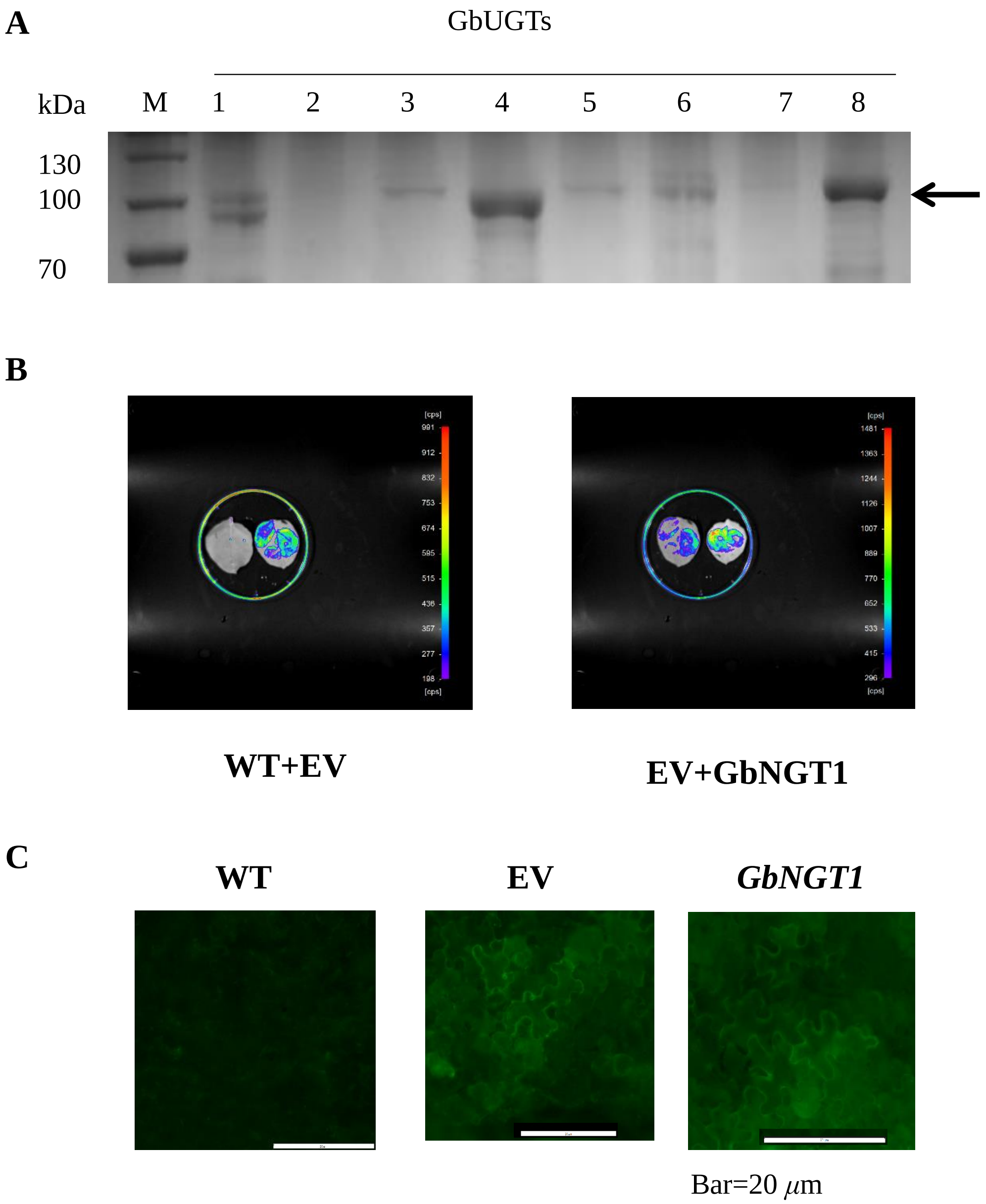


Fig. S3. GbNGT1 expressed in *E.coli* and *N. benthamiana*.

*(A)* SDS-PAGE gel of recombinant GbNGT1 protein; *(B)* and *(C)* transient expressed GFP or *GbNGT1*-GFP in tobacco was confirmed by LB 985 NightShade (Berthold Techonologies) and OLYMPUS IX73, respectively, WT, wildtype, EV, empty vector.

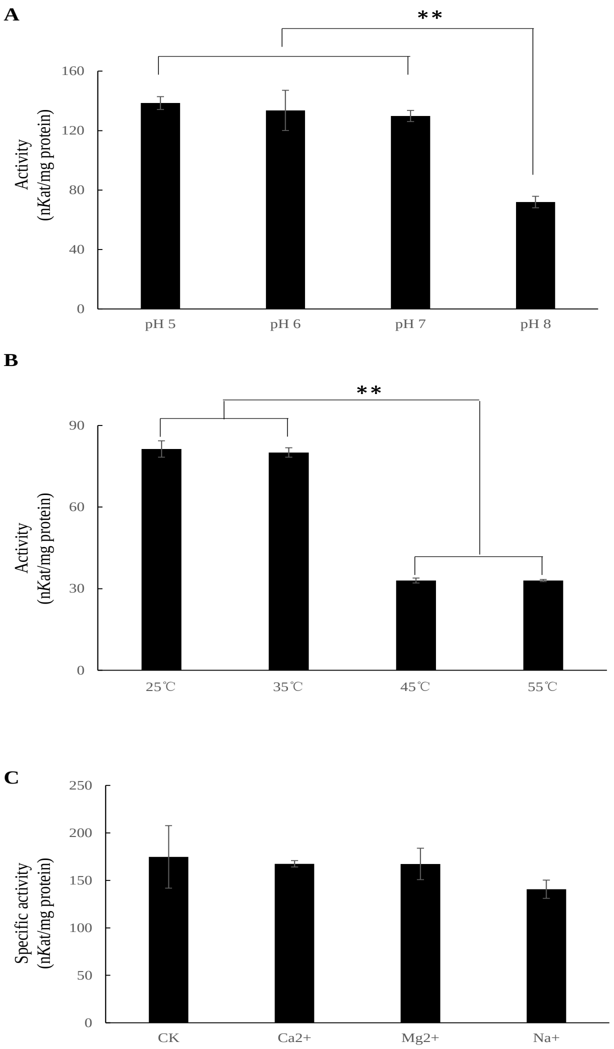


Fig. S4. Characterization of GbNGT1.

The enzymatic activity of GbNGT1 toward IAA-Asp substrate in solution at different pH value *(A)*, temperature *(B)* and metal ion *(C)*.

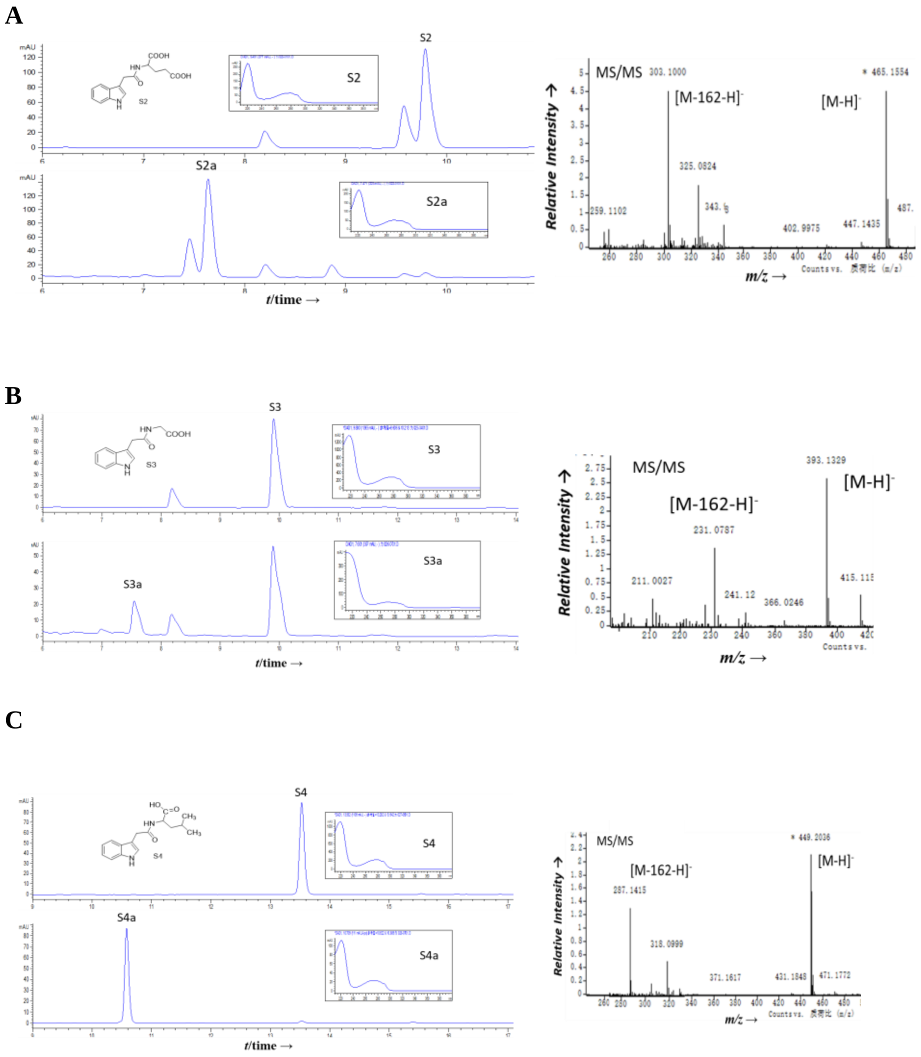


Fig. S5. HPLC and MS spectrums of GbNGT1 with IAA-Glu *(A, S2)*, IAA-Gly *(B, S3)*, and IAA-Leu *(C, S4)*. Negative ion PI model was used to detect the three substrates, S2a, S3a and S4a are the new products in enzymatic reactions.

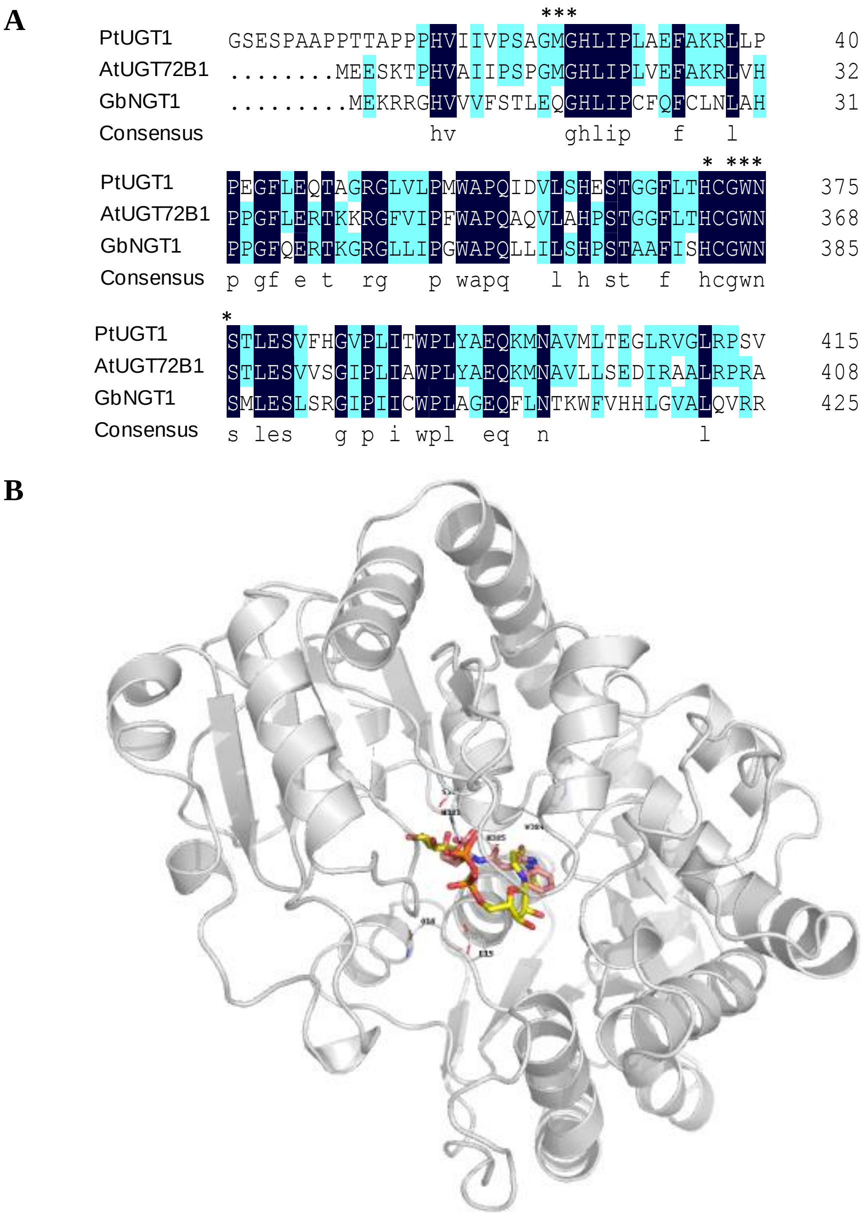


Fig. S6. The key residues predicted by docking.

*(A)* Amino acid alignment of GbNGT1, AtUGT72B1 and PtUGT1, asterisks for binging amino acids. *(B)* The overall chart of GbNGT1 docking with UDPG and IAA-Asp, the molecule marked yellow and orange is UDPG that marked pink is IAA-Asp.

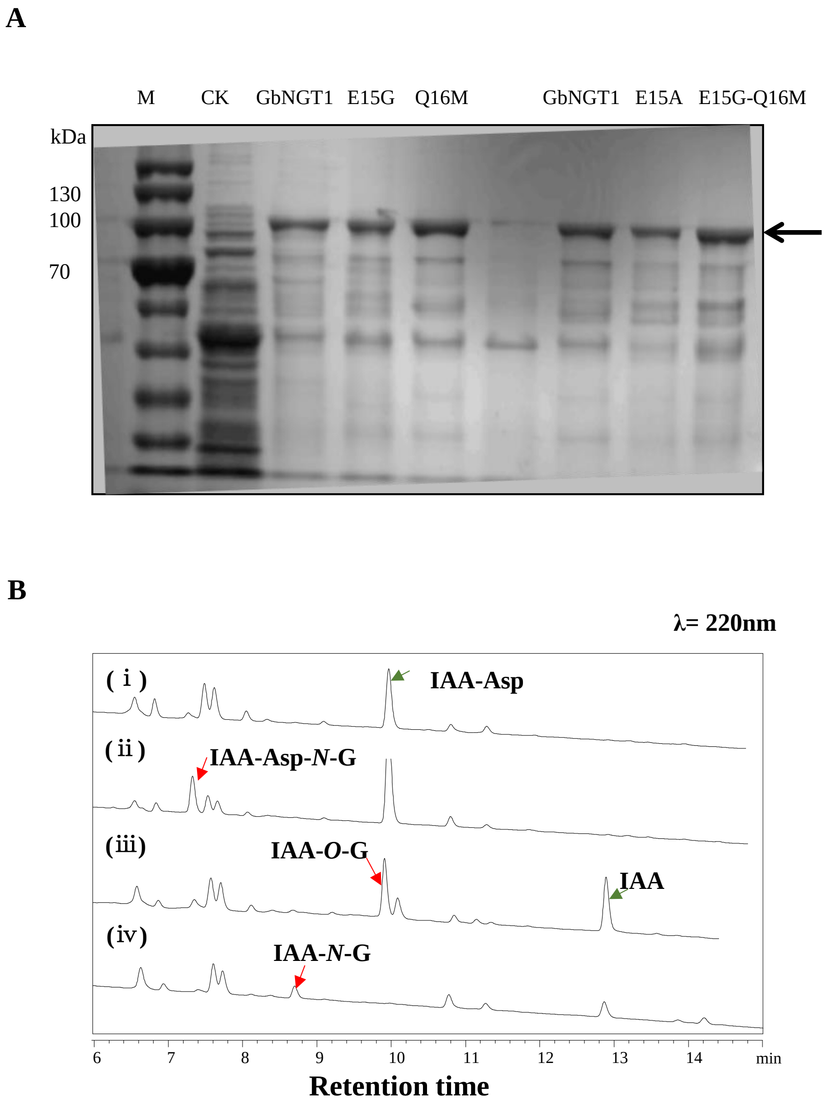


Fig. S7. The functions of GbNGT1 mutants in *E. coli* and *N. benthamiana*.

(A) The SDS-PAGE gel of native and mutant recombinant proteins of GbNGT1. M, maker; CK, empty vector. (B) The HPLC spectrums of *N. benthamiana* leaves transformed mutant *E15G* or *GbNGT1*: (i) mutant *E15G*-transformed leaves adding IAA-Asp; (ii) *GbNGT1*-transformed leaves adding IAA-Asp; (iii) mutant *E15G*-transformed leaves adding IAA; (iv), *GbNGT1*-transformed leaves adding IAA. Red arrow, new product; green arrow, substrate.

Table S1. The 58 ginkgo cultivars used in our study.

| Experimental No. | Names of strains in Chinese | Names of strains in English |
|---|---|---|
| 3 | 铁富 1 号 | No.1, Tiefu |
| 9 | 泰兴 l 号 | No.1, Taixing |
| 10 | 泰兴 2 号 | No.2, Taixing |
| 13 | 洞庭皇 | Dongting Huang |
| 14 | 洞庭佛手 2 | Dongting Foshou 2 |
| 16 | 洞庭佛手 1 | Dongting Foshou 1 |
| 17 | 苏农佛手 | Sunong Foshou |
| 18 | 郯城 231 | Tancheng 231 |
| 19 | 郯城 207 | Tancheng 207 |
| 20 | 延安（1 号）小龙眼 | Yanan (No.1) Xiaolongyan |
| 21 | 港西 2 号 | No.2, Gangxi |
| 23 | 曹楼 1 号 | No.1, Caolou |
| 24 | 曹楼 2 号 | No.2, Caolou |
| 25 | 港西 l 号 | No.1, Gangxi |
| 26 | 铁马 1 号 | No.1, Tiema |
| 27 | 铁马 2 号 | No.2, Tiema |
| 28 | 铁马 3 号 | No.3, Tiema |
| 29 | 铁马 4 号 | No.4, Tiema |
| 30 | 郯城马铃 l 号 （新村花园） | No.1, Tancheng Maling (Xincunhuayuan) |
| 31 | 郯城马铃 2 号 （新村花园） | No.2, Tancheng Maling (Xincunhuayuan) |
| 32 | 大马铃 （新村花园） | Damaling (Xincunhuayuan) |
| 34 | 大龙眼 （安徽全椒） | Dalongyan (Anhui Quanjiao ) |
| 36 | 小园子 | Xiaoyuanzi |
| 37 | 鸭屁股(佛指) | Yapigu (Fo zhi) (Xiongzhu) |
| 39 | 大龙眼 | Dalongyan |

| | | |
|---|---|---|
| 40 | 叶籽银杏 | Yezi Yinxing |
| 43 | 郯城 202 | Tancheng 202 |
| 44 | 新村圆铃 9 号 | No.9, Xincun Yuanling |
| 45 | 新村 18 号 | No.18, Xincun |
| 46 | 贵州正安 l 号 | No.1, Guizhou Zhengan |
| 47 | 贵州正安 2 号 | No.2, Guizhou Zhengan |
| 48 | 贵州正安 3 号 | No.3, Guizhou Zhengan |
| 49 | 贵州正安 4 号 | No.4, Guizhou Zhengan |
| 51 | 道真 5 号 | No.5, Daozhen |
| 52 | 道真 7 号 | No.7, Daozhen |
| 53 | 桂林 2 号 | No.2, Guilin |
| 54 | 桂林 6 号 | No.6, Guilin |
| 56 | 桂林 8 号 | No.8, Guilin |
| 57 | 桂林 9 号 | No.9, Guilin |
| 58 | 浙江长兴 1 号 | No.1, Zhejiang Changxing |
| 59 | 浙江长兴 2 号 | No.2, Zhejiang Changxing |
| 60 | 浙江长兴 3 号 | No.3, Zhejiang Changxing |
| 61 | 浙江长兴 4 号 | No.4, Zhejiang Changxing |
| 63 | 圆铃 13 号 | No.13, Yuanling |
| 65 | 长糯白果 | Changnuo Baiguo |
| 66 | 圆铃 9 号 | No.1, Yuanling |
| 67 | 金坠 5 号 | No.5, Jinzhui |
| 68 | 盘县长白果 | Panxian Changbaiguo |
| 70 | 日本滕久郎 | “Teng Kuo" or "Tengjiulang” |
| 71 | 日本久寿 | "Hisatoshi" or "Jiushou" |
| 72 | 湖北安陆 1-4 | No.1-4, Hubei Anlu |
| 73 | 湖北安陆 1-5 | No.1-5, Hubei Anlu |

| | | |
|---|---|---|
| 75 | 湖北安陆 1-4 | No.1-4, Hubei Anlu |
| 76 | 湖北安陆 1 -6 | No.1-6, Hubei Anlu |
| 78 | 邹庄 6 号 （邳县） | No.6, Zhouzhuang (Pixian) |
| 80 | 如泉(龙眼〉 | Ruquan (Long yan） |
| 82 | 邳县薛集 | Pixian Xueji |
| 83 | 邳县梅核 | Pixian Meihe |

Table S2. The transcripts of 13 candidate GbUGT genes according to published data.

| GbUGTs | Leaf 1 | Leaf2 | Leaf3 | Leaf means | I-fruit1 | I-fruit2 | I-fruit means | R-fruit1 | R-fruit2 | R-fruit means |
|---|---|---|---|---|---|---|---|---|---|---|
| UGT85AH1 | 0.00 | 0.03 | 0.00 | 0.01 | 0.08 | 0.00 | 0.04 | 12.69 | 12.58 | 12.63 |
| UGT74AK2 | 0.15 | 0.42 | 0.05 | 0.21 | 0.35 | 0.00 | 0.18 | 2.77 | 31.65 | 17.21 |
| UGT721B2 | 0.08 | 0.05 | 0.00 | 0.04 | 0.03 | 0.00 | 0.01 | 22.17 | 15.65 | 18.91 |
| UGT85AJ2 | 0.06 | 0.04 | 0.00 | 0.03 | 1.69 | 1.88 | 1.78 | 7.17 | 40.17 | 23.67 |
| UGT85AJ3 | 0.06 | 0.04 | 0.00 | 0.03 | 1.03 | 0.58 | 0.80 | 12.18 | 48.36 | 30.27 |
| UGT721B3 | 0.05 | 0.11 | 0.00 | 0.05 | 0.00 | 0.00 | 0.00 | 41.43 | 28.74 | 35.09 |
| **UGT717A2** **Gb38547** | 1.95 | 1.65 | 0.85 | 1.48 | 6.78 | 6.30 | 6.54 | 38.92 | 36.77 | 37.85 |
| UGT85AJ4 | 0.03 | 0.03 | 0.00 | 0.02 | 0.90 | 0.96 | 0.93 | 219.35 | 197.32 | 208.33 |
| UGT85AJ1 | 31.56 | 20.89 | 35.98 | 29.48 | 0.03 | 0.03 | 0.03 | 2.03 | 2.60 | 2.32 |
| UGT716A1 | 72.08 | 48.27 | 47.70 | 56.02 | 170.12 | 151.51 | 160.81 | 185.78 | 158.81 | 172.29 |
| UGT92K1 | 5.79 | 4.38 | 5.28 | 5.15 | 1.53 | 0.56 | 1.05 | 20.16 | 14.71 | 17.43 |
| UGT725A1 | 23.76 | 5.72 | 25.97 | 18.48 | 17.45 | 20.44 | 18.95 | 0.00 | 0.03 | 0.01 |
| UGT721B1 | 0.38 | 1.15 | 0.20 | 0.58 | 13.90 | 8.53 | 11.21 | 0.15 | 0.58 | 0.36 |
| GbGH3-1/Gb_04369 | 0.00 | 0.00 | 0.00 | 0.00 | 0.48 | 0.09 | 0.28 | 0.00 | 1.73 | 0.87 |
| GbGH3-2/Gb_06148 | 0.02 | 0.00 | 0.08 | 0.03 | 0.13 | 0.33 | 0.23 | 0.03 | 0.04 | 0.04 |
| GbGH3-3/Gb_07002 | 0.07 | 0.06 | 0.00 | 0.04 | 0.11 | 0.00 | 0.06 | 0.00 | 6.79 | 3.39 |
| GbGH3-4/Gb_09255 | 0.01 | 0.00 | 0.00 | 0.00 | 0.04 | 0.00 | 0.02 | 0.16 | 6.69 | 3.43 |
| GbGH3-5/Gb_09499 | 0.19 | 0.07 | 0.19 | 0.15 | 10.19 | 9.92 | 10.05 | 2.37 | 3.10 | 2.74 |
| GbGH3-6/Gb_12334 | 0.14 | 0.00 | 0.00 | 0.05 | 0.01 | 0.00 | 0.01 | 1.99 | 9.75 | 5.87 |
| GbGH3-7/Gb_12335 | 0.55 | 0.04 | 0.79 | 0.46 | 0.01 | 0.22 | 0.12 | 2.86 | 8.70 | 5.78 |
| GbGH3-8/Gb_31649 | 0.00 | 0.00 | 0.00 | 0.00 | 0.06 | 0.00 | 0.03 | 1.20 | 15.59 | 8.39 |
| GbGH3-9/Gb_33150 | 1.80 | 3.53 | 1.68 | 2.34 | 16.76 | 14.97 | 15.86 | 74.22 | 50.89 | 62.56 |

| | | | | | | | | | | |
|---|---|---|---|---|---|---|---|---|---|---|
| GbGH3-10/Gb_36596 | 1.03 | 0.44 | 0.93 | 0.80 | 32.14 | 37.09 | 34.62 | 0.52 | 0.84 | 0.68 |
| GbGH3-11/Gb_41415 | 0.01 | 0.00 | 0.00 | 0.00 | 0.42 | 0.20 | 0.31 | 4.50 | 11.68 | 8.09 |

I-fruit, immature fruit, R-fruit, ripe fruit.

Table S3. The transcripts of 9 cloned GbUGT genes according to transcriptome data collected on June 15th

| Genbank | GbUGTs | Experimental No. | Coat1 | Coat2 | Coat3 | Leaf1 | Leaf2 | Leaf3 | Seed1 | Seed2 | Seed3 |
|---|---|---|---|---|---|---|---|---|---|---|---|
| MN908519 | UGT721B3 | GbUGT1 | 1.02 | 0 | 0 | 0 | 0 | 0 | 0 | 0 | 0 |
| MN908520 | UGT85AJ2 | GbUGT2 | 0.49 | 0 | 0.07 | 0 | 0 | 0 | 0 | 0.08 | 0.18 |
| MN908518 | UGT721B2 | GbUGT3 | 0.69 | 0 | 0 | 0 | 0 | 0 | 1.14 | 0.08 | 1.06 |
| KY274818 | UGT92K1 | GbUGT4 | 1.97 | 0.72 | 0.43 | 1.75 | 2.03 | 2.22 | 1.44 | 0.67 | 0.37 |
| MN908521 | UGT74AK2 | GbUGT5 | 0 | 0 | 0 | 0.24 | 0.16 | 0 | 1.57 | 0 | 0.97 |
| KY274815 | UGT721B1 | GbUGT6 | 41.25 | 84.64 | 37.16 | 0.45 | 0.31 | 0.14 | 4.05 | 7.46 | 0 |
| KY274816 | UGT725A1 | GbUGT7 | 8.56 | 13.28 | 7.97 | 17.82 | 25.7 | 5.52 | 2.79 | 23.97 | 8.19 |
| MN908522 | UGT717A2-Gb38547 | GbUGT8 | 38.14 | 16.63 | 20.33 | 7.94 | 8.65 | 5.39 | 50.29 | 41.58 | 61.53 |
| KX371617 | UGT716A1 | GbUGT9 | 152.2 | 233.79 | 115.38 | 60.39 | 32.37 | 27.2 | 47.47 | 91.18 | 61.68 |
| MN908516 | / | GbGH3.2 | 0.41 | 0.07 | 0.22 | 3.06 | 0.3 | 0 | 1.24 | 2.3 | 0.72 |
| MN908517 | / | GbGH3.5 | 48.7 | 3.44 | 9.8 | 0.48 | 0.24 | 0 | 66.95 | 96.87 | 97.69 |

Table S4 [1]H-NMR and [13]C-NMR spectrum data of IAA-Asp-*N*-Glc compounds

| 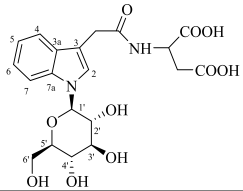 | IAA-Asp-*N*-Glc | |
|---|---|---|
| | $\delta_C$ | $\delta_H$ |
| 2 | 124.3 | 7.40 (1H, *t*) |
| 3 | 109.2 | |
| 3a | 137.1 | - |
| 4 | 118.4 | 7.54 (2H, *t*) |
| 5 | 119.6 | 7.10 (1H, *t*) |
| 6 | 121.8 | 7.19 (1H, *t*) |
| 7 | 110.1 | 7.54 (2H, *t*) |
| 7a | 128.3 | - |
| $-CH_2CONH$ | 32.1 | 3.74 (2H, *m*) |
| -C=O | 172.8 | - |
| α | 49.1 | 4.77 (1H, *m*) |
| β | 35.5 | 2.84 (2H, *m*) |
| 1’ | 85.3 | 5.44 (1H, *d*, *J*=12Hz) |
| 2’ | 72.4 | 3.93 (1H, *m*) |
| 3’ | 77.4 | 3.63 (1H, *m*) |
| 4’ | 79.1 | 3.61 (1H, *m*) |
| 5’ | 70.1 | 3.53 (1H, *m*) |
| 6’ | 61.3 | 3.90 (1H, *d*, *J*=6 Hz)<br>3.74 (1H, *d*, *J*=12 Hz) |
| -COOH | 172.9 | |
| -COOH | 172.9 | |

Table S5 $^1$H-NMR and $^{13}$C-NMR spectrum data of IAA-Gly-*N*-Glc compounds

| 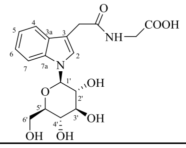 | IAA-Gly-*N*-Glc | |
|---|---|---|
| | $\delta_C$ | $\delta_H$ |
| 2 | 124.9 | 7.42 (1H, *t*) |
| 3 | 109.9 | - |
| 3a | 136.6 | - |
| 4 | 119.6 | 7.54 (2H, *t*) |
| 5 | 120.7 | 7.18 (1H, *t*) |
| 6 | 122.8 | 7.27 (1H, *t*) |
| 7 | 110.3 | 7.54 (2H, *t*) |
| 7a | 127.8 | - |
| -$CH_2$CONH | 31.9 | 3.75 (2H, *m*) |
| -C=O | 174.9 | - |
| α | 42.5 | 3.74 (2H, *m*) |
| 1’ | 84.3 | 5.56 (1H, *d*, *J*=12Hz) |
| 2’ | 71.6 | 4.01 (1H, *m*) |
| 3’ | 78.3 | 3.67 (1H, *m*) |
| 4’ | 76.4 | 3.60 (1H, *m*) |
| 5’ | 69.3 | 3.58 (1H, *m*) |
| 6’ | 60.5 | 3.82 (1H, *d*, *J*=12Hz) |
| | | 3.68 (1H, *d*, *J*=8Hz) |
| -COOH | 175.9 | - |

Table S6 $^{1}H$-NMR and $^{13}C$-NMR spectrum data of IAA-Leu-*N*-Glc compounds

| 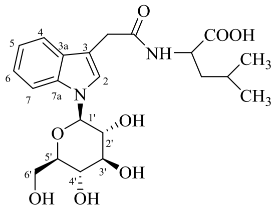 | IAA-Leu-*N*-Glc | |
|---|---|---|
| | $\delta_C$ | $\delta_H$ |
| 2 | 124.7 | 7.40 (1H, *t*) |
| 3 | 109.9 | - |
| 3a | 136.7 | - |
| 4 | 119.1 | 7.54 (2H, *t*) |
| 5 | 120.7 | 7.18 (1H, *t*) |
| 6 | 122.9 | 7.26 (1H, *t*) |
| 7 | 110.4 | 7.54 (2H, *t*) |
| 7a | 127.8 | - |
| -$CH_2$CONH | 32.1 | 3.72 (2H, *m*) |
| -C=O | 174.7 | - |
| α | 52.1 | 4.29 (1H, *m*) |
| β | 39.4 | 1.53 (2H, *m*) |
| γ | 24.4 | 1.44 (1H, *m*) |
| 1’ | 84.3 | 5.55 (1H, *d*, *J*=12Hz) |
| 2’ | 71.7 | 3.98 (1H, *m*) |
| 3’ | 78.3 | 3.67 (1H, *m*) |
| 4’ | 76.4 | 3.60 (1H, *m*) |
| 5’ | 69.3 | 3.58 (1H, *m*) |
| 6’ | 60.5 | 3.82 (1H, *d*, *J*=12Hz) |
| | | 3.68 (1H, *d*, *J*=8Hz) |
| -$CH_3$ | 20.3 | 0.73 (3H, *d*, *J*=6Hz) |
| -$CH_3$ | 22.2 | 0.77 (3H, *d*, *J*=6Hz) |
| -COOH | 177.4 | |

Table S7 The predicted docking energy of GbNGT1 with IAA and IAA-AAs.

| Name | Total Energy | VDW | HBond | Elec | AverConPair |
|---|---|---|---|---|---|
| IAA | -101.119 | -82.6159 | -15.4375 | -3.06515 | 47.3846 |
| IAA-Asp | -120.568 | -87.7991 | -32.4626 | -0.306465 | 31.2381 |
| IAA-Gly | -108.639 | -84.989 | -22.6968 | -0.953459 | 28.5909 |
| IAA-Glu | -124.958 | -94.6143 | -30.8444 | 0.500643 | 30.2727 |
| IAA-Leu | -113.638 | -94.8505 | -18.8403 | 0.0530466 | 30.7143 |

The output data included total energy (Kcal/mol), van der Waals interactions (VDW, Kcal/mol), Hydrogen bonding (HBond, Kcal/mol), electrostatic interactions (Elec Kcal/mol), and average conpair (AverConPair).

**Appendix S** Synthesis of Substrates and NMR information for used compounds

Synthesis of Substrates

The synthesis steps mainly include following procedure: Firstly, methylation reactions of L-aspartic acid (Asp, aspartic acid), L-glutamic acid (Glu, glutamic acid), glycine (Gly, glycine) and L-leucine (Leu, Leucine) were performed. Taking the Asp methylation reaction as an example, 3 g of Asp (22.6 mmol) was taken into a 100 mL reaction flask, and 31 mL of methanol was added into, then the reaction flask was placed on a magnetic stirrer. 2 mL of $SOCl_2$ (27 mmol, 1.2 equivalents) was added dropwise at a low temperature. After dissolving aspartic acid completely, the reaction flask was placed in a 100 ℃ oil bath and refluxed for 4 h, then the reaction was stopped, and this solvent was recovered to a residue on a vacuum rotary evaporator. According to above mentioned method, others three amino acids were methylated. The reaction equation was shown as following:

a

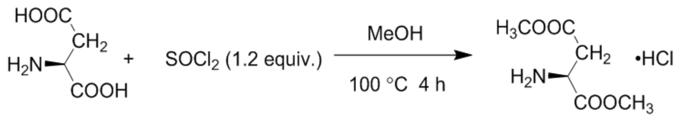


b

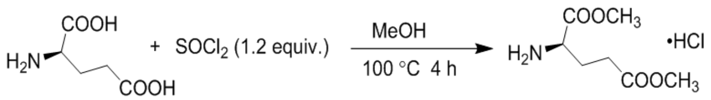


c

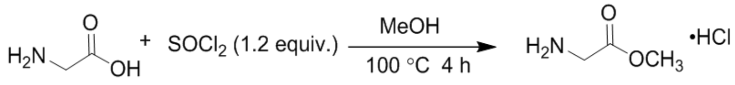


d

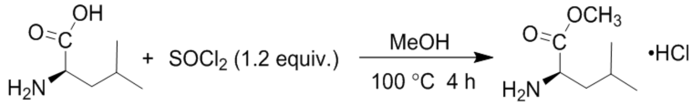


Secondly, amidation reaction was conducted; indoleacetic acid (IAA), 5-methylindoleacetic acid (5-Me-IAA), 5-Br indoleacetic acid (5-Br-IAA) or indole-2-acetic acid (2-IAA) were synthesized using the above obtained products, including dimethyl aspartate (Asp-(OMe) -OMe), dimethyl glutamate (Glu-(OMe) -OMe), glycine methyl ester (Gly-OMe), and leucine methyl ester (Leu -OMe) or Propylamine. Taking the amidation reaction of IAA and Asp (OMe)-OMe as an instance, IAA (300 mg, or 1.7 mmol), Asp (OMe) -OMe (2.5 mmol, 1.5 equivalent), 1.2 folds equivalent N, N'- dicyclohexylcarbodiimide (DCC, 2 mmol), 0.2 folds equivalent of 4-dimethylaminopyridine (DMAP, 0.34 mmol), 2.5 folds equivalent of N, N-diisopropylethylamine (DIPEA, 4.3 mmol) and acetonitrile (8 mL) were mixed in a 25 mL reaction tube; the reaction was performed on a magnetic stirrer at room temperature. Using IAA as a reference substance, the reaction solution was analyzed by thin layer chromatography. The product was isolated from the reaction solution after the IAA was almost consumed completely; it was identified by NMR finally.

Eventually, the amidation products were hydrolyzed, except for the amidation products of IAA. Taking the hydrolysis reaction of IAA-Asp-(OMe)-OMe as an example, IAA-Asp-(OMe)-OMe was dissolved in NaOH $H_2O$ (2M)and EtOH $H_2O$ ($V_{EtOH}$ :$V_{H2O}$ =5:5), then placed in a 25 mL reaction tube, and reacted on a magnetic stirrer at room temperature. Using IAA-Asp-(OMe)-OMe as a reference, the reaction solution was analyzed by thin layer chromatography. The reaction was finished after the IAA-Asp-(OMe)-OMe reaction was almost complete. After recovering the solvent, water was dissolved and extracted with ethyl acetate. Consequently, ethyl acetate was recovered to obtain targeted products.

The NMR data of each compound are as following:

**IAA-Asp (OMe)-OMe:** As orange-yellow oil, $^{13}$C-NMR ($CDCl_3$, 150 MHz): 171.7 (C-9), 171.1 (C-5'), 171.1 (C-4'), 136.6 (C-7a), 126.9 (C-3a), 124.1 (C-2), 122.1 (C-6), 119.5 (C-4), 118.4 (C-5), 111.6 (C-7), 107.8 (C-3), 52.7 (C-2'), 51.8 (C-7'), 48.5 (C-6'), 36.0 (C-3'), 33.3 (C-8).

**IAA-Asp (1):** As yellow-white solid, $^{1-3}$C-NMR ($CD_3OD$, 150 MHz): 174.2 (C-5'), 174.0 (C-4'), 173.2 (C-9), 138.1 (C-7a), 128.5 (C-3a), 125.2 (C-2), 122.7 (C-6), 120.1 (C-4), 119.5 (C-5), 112.4 (C-7), 108.9 (C-3), 50.3 (C-2'), 36.8 (C-3'), 33.8 (C-8).

**5-Me-IAA-Asp (OMe)-OMe (intermediate product):** As white solid, $^{13}$C-NMR ($CD_3OD$, 150 MHz): 175.2 (C-9), 173.0 (C-5'), 172.8 (C-4'), 137.0 (C-7a), 129.6 (C-3a), 129.1 (C-5), 125.7 (C-2), 124.8 (C-6), 119.5 (C-4), 112.6 (C-7), 108.9 (C-3), 53.5 (C-2'), 52.8 (C-7'), 50.7 (C-6'), 34.3 (C-3'), 31.2 (C-8), 22.3 (C-10).

**5-Me-IAA-Asp (5):** As powder-white solid, $^{13}$C-NMR ($CD_3OD$, 150 MHz): 174.9 (C-5'), 174.1 (C-4'), 174.0 (C-9), 136.6 (C-7a), 129.3 (C-3a), 128.9 (C-5), 125.3 (C-2), 124.4 (C-6), 119.2 (C-4), 112.1 (C-7), 108.5 (C-3), 50.3 (C-2'), 36.9 (C-3'), 33.9 (C-8), 21.8 (C-10)

**5-Br-IAA-Asp (OMe)-OMe (intermediate product):** As yellow oil, $^{13}$C-NMR ($CDCl_3$, 150 MHz): 171.6 (C-9), 171.4 (C-5'), 171.3 (C-4'), 135.3 (C-7a), 128.8 (C-3a), 125.5 (C-6), 125.1 (C-2), 121.2 (C-4), 113.2 (C-7), 113.0 (C-5), 107.7 (C-3), 53.0 (C-2'), 52.2 (C-7'), 48.7 (C-6'), 36.1 (C-3'), 33.3 (C-8).

**5-Br-IAA-Asp (9):** As yellow solid, $^{13}$C-NMR ($CD_3OD$, 150 MHz): 174.5 (C-5'), 174.1 (C-4'), 174.1 (C-9), 136.9 (C-7a), 130.5 (C-3a), 126.7 (C-6), 125.4 (C-2), 122.3 (C-4), 114.1 (C-7), 113.3 (C-5), 109.1 (C-3), 50.4 (C-2'), 36.9 (C-3'), 33.6 (C-8).

**IAA-Glu (OMe)-OMe**: As yellow oil, $^{13}$C-NMR ($CDCl_3$, 150 MHz): 173.0 (C-5'), 172.2 (C-6'), 172.0 (C-9), 136.5 (C-7a), 126.9 (C-3a), 124.1 (C-2),122.0 (C-6), 119.4 (C-4),118.3 (C-5),111.5 (C-7), 107.7 (C-3), 53.5 (C-2'), 52.2 (C-8'), 51.6 (C-7'), 33.3 (C-8), 29.8 (C-3'), 26.8 (C-4').

**IAA-Glu (2):** As orange-red solid, $^{13}$C-NMR ($CD_3OD$, 150 MHz): 176.5 (C-5'), 175.2 (C-6'), 175.0 (C-9), 138.2 (C-7a), 128.6 (C-3a), 125.1 (C-2), 122.7 (C-6), 120.1 (C-4), 119.5 (C-5), 112.5 (C-7), 109.3 (C-3), 53.2 (C-2'), 33.9 (C-8), 31.2 (C-3'), 28.0 (C-4').

**5-Me-IAA-Glu (OMe)-OMe (intermediate product):** As yellow oil, $^{13}$C-NMR ($CDCl_3$, 150 MHz): 173.2 (C-5'), 172.3 (C-6'), 172.0 (C-9), 135.0 (C-7a), 129.3 (C-5), 127.4 (C-3a), 124.3 (C-2), 124.1 (C-6), 118.3 (C-4), 111.3 (C-7), 107.9 (C-3), 52.6 (C-2'), 51.9 (C-7), 51.7 (C-7'), 33.5 (C-8), 30.0 (C-3'), 27.2 (C-4'), 21.6 (C-10).

**5-Me-IAA-Glu (6):** As brown-red solid, $^{13}$C-NMR ($CD_3OD$, 150 MHz): 176.4 (C-5'), 175.3 (C-6'), 175.0 (C-9), 136.6 (C-7a), 129.3 (C-5), 128.8 (C-3a), 125.3 (C-2), 124.4 (C-6), 119.2 (C-4), 112.2 (C-7), 108.7 (C-3), 53.0 (C-2'), 34.0 (C-8), 31.0 (C-3'), 27.9 (C-4'), 21.8 (C-10).

**5-Br-IAA-Glu (OMe)-OMe (intermediate product):** As yellow solid, $^{13}$C-NMR ($CD_3OD$, 150 MHz): 174.9 (C-5'),174.8 (C-6'), 173.7 (C-9), 136.9 (C-7a),130.5 (C-3a),126.7 (C-6), 125.4 (C-2),122.3 (C-4), 114.1 (C-5), 113.3 (C-7),109.3 (C-3),53.3 (C-2'), 53.0 (C-8'), 52.3 (C-7'), 33.7 (C-8), 31.0 (C-3'), 27.5(C-4').

**5-Br-IAA-Glu (10):** As orange-yellow solid, $^{13}$C-NMR ($CD_3OD$, 150 MHz): 176.5 (C-5'), 174.9 (C-6'), 174.9 (C-9), 136.9 (C-7a), 130.5 (C-3a), 126.6 (C-6), 125.4 (C-2), 122.3 (C-4), 114.1 (C-5), 113.3 (C-7), 109.4 (C-3), 53.3 (C-2'), 33.6 (C-8), 31.3 (C-3'), 28.0 (C-4').

**IAA-Gly (3):** As light-yellow solid, $^{13}$C-NMR ($CD_3OD$, 150 MHz): 177.1 (C-3'), 174.8 (C-9), 138.1 (C-7a), 128.6 (C-3a), 125.3 (C-2), 122.6 (C-6), 120.0 (C-5), 119.4 (C-4), 112.5 (C-7), 109.3 (C-3), 50.0 (C-2'), 23.8 (C-8).

**5-Me-IAA-Gly-OMe (intermediate product):** As white solid, $^{13}$C-NMR ($CD_3OD$, 150 MHz): 175.7 (C-9), 171.9 (C-3'), 136.6 (C-7a), 129.2 (C-5), 128.9 (C-3a), 125.3 (C-2), 124.4 (C-6), 119.2 (C-4), 112.2 (C-7), 108.6 (C-3), 52.7 (C-4'), 42.2 (C-2'), 33.8 (C-8), 21.8 (C-10).

**5-Me-IAA-Gly (7) :** As orange-yellow solid, $^{13}$C-NMR ($CD_3OD$, 150 MHz): 175.7 (C-3'), 173.1 (C-9), 136.6 (C-7a), 129.3 (C-5), 128.9 (C-3a), 125.3 (C-2), 124.4 (C-6), 119.2 (C-4), 112.2 (C-7), 108.6 (C-3), 42.1 (C-2'), 33.8 (C-8), 21.8 (C-10).

**5-Br-IAA-Gly-OMe (intermediate product):** As light-green oil, $^{13}$C-NMR ($CDCl_3$, 150 MHz): 172.1 (C-9), 170.5 (C-3'), 135.2 (C-7a), 128.9 (C-3a), 125.5 (C-6), 125.4 (C-2), 121.3 (C-4), 113.2 (C-5), 133.2 (C-7), 107.9 (C-3), 52.5 (C-4'), 41.5 (C-2'), 33.0 (C-8).

**5-Br-IAA-Gly (11):** As yellow solid, $^{13}$C-NMR ($CD_3OD$, 150 MHz): 175.1 (C-9), 173.1 (C-3'), 136.8 (C-7a), 130.5 (C-3a), 126.7 (C-6), 125.4 (C-2), 122.3 (C-4), 114.1 (C-5), 113.4 (C-7), 109.2 (C-3), 42.1 (C-2'), 33.6 (C-8).

**IAA-Leu-OMe (intermediate product):** As orange-yellow oil, $^{13}$C-NMR ($CDCl_3$, 150 MHz): 173.5 (C-6'), 171.7 (C-9), 136.7 (C-7a), 127.2 (C-3a), 124.0 (C-2), 122.7 (C-6), 120.1 (C-5), 118.9 (C-4), 111.5 (C-7), 108.7 (C-3), 52.4 (C-2'), 50.9 (C-8'), 41.4 (C-3'), 33.3 (C-8), 24.8 (C-4'), 22.9 (C-5'), 22.0 (C-7').

**IAA-Leu (4):** As powder-white solid, $^{13}$C-NMR ($CD_3OD$, 150 MHz)： 176.2 (C-6'), 175.0 (C-9), 138.3 (C-7a), 128.7 (C-7a), 125.1 (C-2), 122.7 (C-6), 120.0 (C-5), 119.6 (C-4), 112.4 (C-7), 109.5 (C-3), 52.3 (C-2'), 41.8 (C-3'), 33.9 (C-8), 26.1 (C-4'), 23.5 (C-5'), 21.9 (C-7').

**5-Me-IAA-Leu-OMe (intermediate product):** As yellow oil, $^{13}$C-NMR ($CD_3OD$, 150 MHz)： 175.2 (C-6'), 174.8 (C-9), 136.6 (C-7a), 129.1 (C-5), 128.8 (C-3a), 125.2 (C-2), 124.3 (C-5), 119.3 (C-4), 112.2 (C-7), 108.9 (C-3), 52.7 (C-2'), 52.4 (C-8'), 41.5 (C-3'), 33.9 (C-8), 26.0 (C-4'), 23.5 (C-10), 21.8 (C-5', 7').

**5-Me-IAA-Leu (8):** As orange solid，$^{13}$C-NMR ($CD_3OD$, 150 MHz)：176.1 (C-6'), 175.1 (C-9), 136.6 (C-7a), 129.1 (C-5), 128.8 (C-3a), 125.2 (C-2), 124.3 (C-5), 119.2 (C-4), 112.2 (C-7), 108.9 (C-3), 52.2 (C-2'), 41.8 (C-3'), 34.0 (C-8), 26.2 (C-4'), 23.5 (C-10), 21.9 (C-5'), 21.9 (C-7').

**5-Br-IAA-Leu-OMe (intermediate product):** As yellow-white solid, $^{13}$C-NMR ($CD_3OD$, 150 MHz)：174.8 (C-6'), 174.7 (C-9), 136.9 (C-7a), 130.4 (C-3a), 126.6 (C-6), 125.4 (C-2), 122.3 (C-4), 114.1 (C-5), 113.2 (C-7), 109.4 (C-3), 52.8 (C-2'), 52.4 (C-8'), 41.5 (C-3'), 33.7 (C-8), 26.1 (C-4'), 23.5 (C-5'), 21.8 (C-7').

**5-Br-IAA-Leu (12):** As orange-yellow solid, $^{13}$C-NMR ($CD_3OD$, 150 MHz)：176.1 (C-6'), 174.7 (C-9), 136.8 (C-7a), 130.4 (C-3a), 126.6 (C-6), 125.4 (C-2), 122.3 (C-4), 114.1 (C-5), 113.2 (C-7), 109.4 (C-3), 52.3 (C-2'), 41.7 (C-3'), 33.7 (C-8), 26.1 (C-4'), 23.6 (C-5'), 21.8 (C-7').
